# Altered Crosstalk of Bacterial Lipopolysaccharide with Immune Cells in Colorectal Cancer Compared to Paired Adjacent Intestinal Tissue

**DOI:** 10.64898/2026.02.01.701966

**Authors:** Åsa Walberg, Anna Maria Reuss, Reihane Ziadlou, Céline Mamie, Claudia Gottier, Anna White, Mohammadmilad Ameri, Marie-Charlotte Brüggen, Matthias Turina, Michaela Ramser, Paulina Wawrzyniak, Maria Walker, Luca Truscello, Adriano Aguzzi, Anne Müller, Barbara Hubeli, Yasser Morsy, Michael Scharl

**Affiliations:** Department of Gastroenterology and Hepatology, University Hospital Zurich, University of Zurich, Zurich, Switzerland; Institute of Neuropathology, University Hospital Zurich, University of Zurich, and ZNZ Neuroscience Center, Zurich, Switzerland; Department of Dermatology, University of Zurich, Zurich, Switzerland; Department of Visceral and Transplant Surgery, University Hospital Zurich, Zurich, Switzerland; Institute of Molecular Cancer Research, University of Zurich, Zurich, Switzerland; Comprehensive Cancer Center Zurich, University Hospital Zurich, University of Zurich; Zurich, Switzerland

**Keywords:** Gut microbiota, colorectal cancer, bacterial-immune interactions, bacterial lipopolysaccharide (LPS), tumor microenvironment, spatial transcriptomics, imaging mass cytometry, 3D imaging iDISCO

## Abstract

Commensal bacteria play a crucial role in modulating human immune responses in the intestine. Under homeostatic conditions, gut microbiota are tightly regulated by interactions with the mucosal immune system. However, colorectal cancer (CRC) is characterized by an imbalance in bacterial composition and bacterial translocation across the intestinal barrier. The spatial distribution of bacteria and their interactions with immune cells in CRC tumors are poorly understood. By applying 3D light-sheet imaging, spatial transcriptomics, and imaging mass cytometry to patient-derived CRC and adjacent tissue, bacterial lipopolysaccharide (LPS) is visualized alongside immune cells and vessels. The results show regional bacterial LPS accumulation and colocalization with distinct immune cell subsets. In CRC-adjacent tissue, bacterial LPS is mainly associated with CD11c^+^ dendritic cells, CD15^+^ neutrophils, and CD163^+^ macrophages. In matched CRC tissue, the number and LPS colocalization of CD163^+^ macrophages and CD11c^+^ dendritic cells decreased, while CD15^+^ neutrophils and their colocalization with LPS increased. Notably, immune cell composition and immune cell-bacteria interactions differ between tumor and adjacent tissue, offering insights into host-microbiota dynamics and mechanistic interactions.

## INTRODUCTION

The human gut microbiome, as well as intra-tumoral bacteria, are pivotal in the development and progression of cancer^1-7^. Microbiota-immune cell interactions in the intestine require both tolerance for commensal bacteria as well as defense mechanisms against harmful pathogens. Under homeostatic conditions, several defense mechanisms control and compartmentalize the gut bacteria to the mucus layer that covers the intestinal epithelium^8^. In addition to epithelial and mucosal barriers, both innate and adaptive immune mechanisms contribute to the regulation of the microbial composition. For instance, dendritic cells (DCs) continuously sample and present bacterial antigens to stimulate antigen-specific T cell responses. Goblet cells in the epithelium can transcytose gut luminal contents, including bacterial antigens, resulting in T cell-mediated antigen-specific tolerance^9^. In contrast, B-/plasma cells contribute to intestinal homeostasis through the production of secretory IgA^10^.

However, in cancer patients, the gut microbiome is frequently characterized by alterations in bacterial composition and function^11^. These alterations impact on gut barrier and immune cell responses^11-13^. Furthermore, certain bacterial species were found in particularly large numbers within the colorectal cancer (CRC) tissue^3,14,15^. Though it remains unclear, how bacteria infiltrate cancerous tissues, some reports suggest that the CRC-associated bacterium, *Fusobacterium nucleatum*, can translocate via a hematological route to the CRC tumors^16,17^. Particularly in CRC, an apparent origin of bacteria detectable within the tumor microenvironment (TME) is the gut lumen, from where bacterial translocation likely originates^18^. Alterations in the TME and disruption of the epithelial and mucosal barrier, as typically found in CRC, may favor the enrichment of selected bacteria at tumor sites^19^. Of note, both pro-tumorigenic and anti-tumorigenic effects of bacteria have been shown in animal models and human studies^5,20,21^. However, the specific bacteria-immune interactions in tumor and tumor-adjacent tissue, as well as the spatial location of bacteria in tumors, are not well characterized. Bacteria have been described mainly intracellularly in cancer cells and immune cells within tumor tissue, but also extracellularly in cancer tissue of various tumor types^22-25^. In CRC, biofilm formations, i.e., large aggregates of bacteria have been found in the unaffected adjacent colon as well as in the CRC tissue^3,26-28^.

In our present study, we analyzed paired CRC tissue and adjacent intestinal tissue from the same patient. We focused on bacterial lipopolysaccharide (LPS) as a general marker for gram-negative bacteria, such as *Fusobacterium nucleatum* or *Bacteroides fragilis*, to assess localized alterations in microbiota-immune interactions in patients with CRC. A better characterization and particularly understanding of bacteria-immune cell interactions in patient-derived tissue will not only help to improve our understanding of the regions populated by bacteria in CRC tumors, but also provide important insights into the mechanistic effects of bacteria on CRC pathogenesis, which in turn could contribute to our understanding of bacteria-based cancer therapies.

## Results

### 3D imaging demonstrates aggregates of bacterial LPS and colocalization with immune cells in CRC tissue

To obtain a global overview of bacterial and immune cell localization within the CRC adjacent intestinal tissue and the CRC tissue, we applied organic solvent-based tissue clearing and immunolabeling of approximately 5 mm^3^-sized patient-derived tissue samples. We applied bacterial LPS as a marker for gram-negative bacteria, CD45 as a general immune cell marker, and PVALP as a marker for the vasculature. In total, six patients with CRC were included in the analysis, comprising six adjacent CRC intestinal tissue samples and five matched CRC samples from the same patients. In the CRC adjacent intestinal tissue, bacterial LPS was detected mainly in the epithelial cell layer, likely deriving from bacteria-rich mucus remnants (Fig. 1A-C, G, Suppl. Video 1). Additionally, in 3/6 CRC adjacent intestinal tissue samples, we observed bacterial LPS translocation into subepithelial/submucosal layers (Fig. S1). In contrast, in CRC tissues, we observed selective regions of LPS, indicative of bacterial abundance, rather than an even distribution across the whole tissue (Fig. 1D-F, H, Suppl. Video 2). The comparison between the matched CRC tissues and the CRC adjacent intestinal tissues revealed an increase in bacterial LPS per volume (mm^3^) in 4/5 CRC tissues (Fig. 1I). However, the extent of increase varied substantially between patients, indicating high inter-patient variability in LPS accumulation. Of note, only a few of the bacterial LPS signals were colocalized with blood vessels (Fig. 1J). The mean colocalization of bacterial LPS with PVALP, indicative of blood vessels, was 0.3 ± 0.05% SD in CRC tissue and slightly lower (0.2 ± 0.07% SD) in CRC adjacent intestinal tissue. Additionally, our results indicated a higher number of blood vessels per mm^3^ in the tumor tissue (5817± 2839 SD) than in the CRC adjacent intestinal tissues (2602 ±1163 SD) (Fig. S2A, B). Across all samples, we obtained high percentages of colocalization of bacterial LPS with CD45+ immune cells (Fig. 1K, Fig. S2C, D). The mean of LPS–CD45 colocalization was 90% in CRC adjacent intestinal tissue samples and 80% in CRC tissue samples. Overall, our 3D staining revealed an increase in bacterial LPS within the CRC tissue, co-localizing with immune cells.

**Figure 1:**
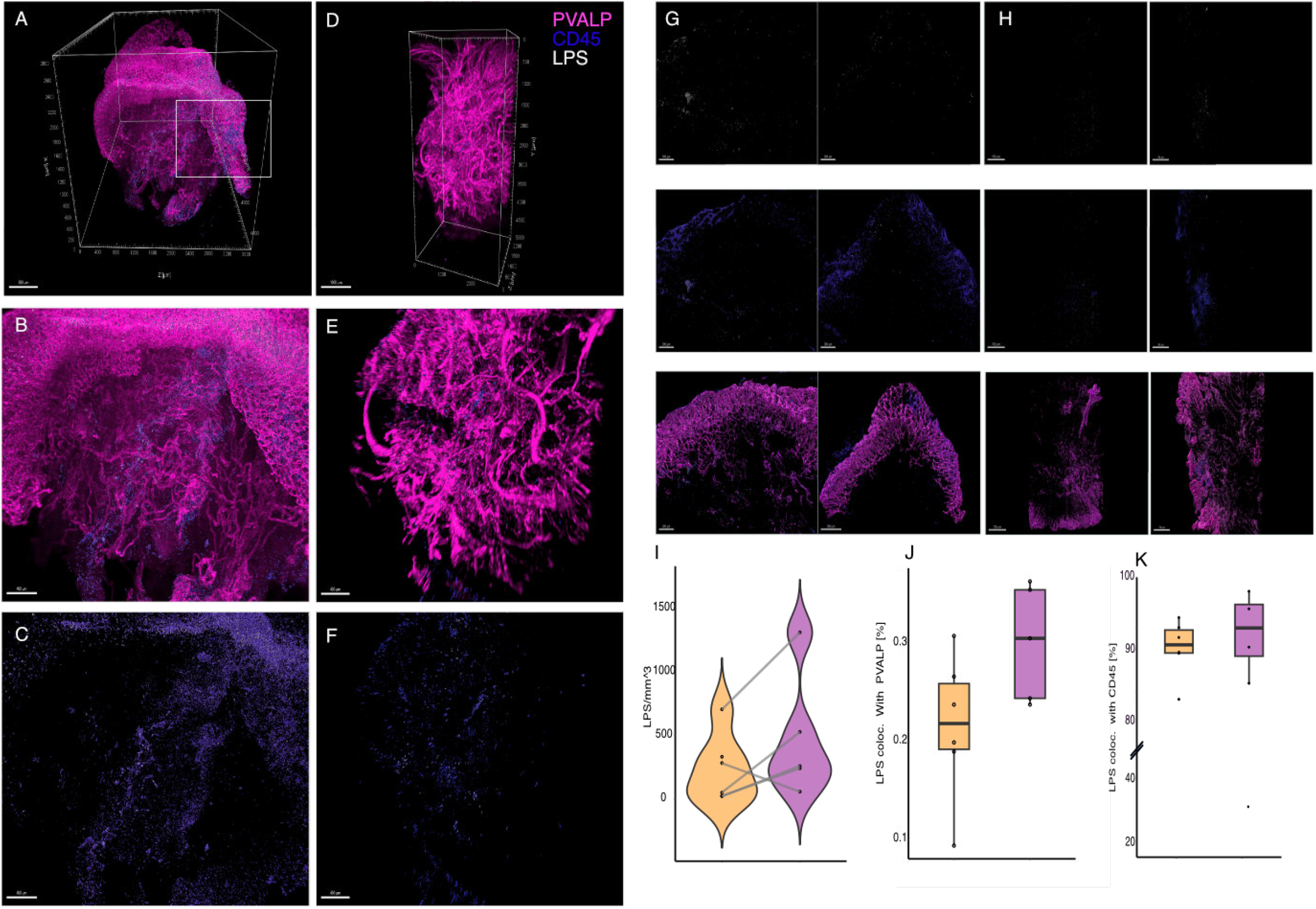
Global overview of bacteria-immune cell interactions in CRC adjacent intestinal tissue and CRC tissue using 3D histology. (**A**): Volume rendering of vasculature (PVALP), immune cells (CD45), and bacterial lipopolysaccharide (LPS) in CRC adjacent intestinal tissue, from xz-, yz-, and xy-views and with zoom-ins**(B)** (white boxes), and Segmented LPS, CD45 objects are displayed **(C)**. (**D):** Volume rendering of CRC tissue stained with the same markers as described in (A), with zoom-ins **(E)** and Segmented LPS, CD45 objects are displayed **(F). (G)** adjacent intestinal tissue from two different CRC samples. Segmented LPS, CD45, and PVALP objects are displayed, and CRC tissue **(H). (I):** Violin plot presenting the ratio of detected bacterial LPS per mm^3^ in CRC adjacent intestinal tissue (orange) and CRC tissue (violet). Lines connect the CRC adjacent intestinal tissue and the CRC tissue from the same patient. **(J-K):** Percentage of bacterial LPS colocalized with the vasculature (J) and immune cells (K) in CRC adjacent intestinal tissue(orange) and CRC tissue (violet). Boxes represent the interquartile range, with the median indicated by a horizontal line. n=6 CRC adjacent intestinal tissue samples and n=5 matched tumor samples.

### Spatial transcriptomics analysis detect inflamed epithelial cells as increased in paired CRC tissue compared to adjacent intestinal tissue

To gain an overview of the immune cell landscape in CRC tissue and paired adjacent tissue from the same patients, we next applied spatial transcriptomics on tissues from three patients with CRC. We detected 10 different cluster cell types within the spatial transcriptomics data of both CRC-adjacent intestinal tissue and CRC tissue (Fig. 2A, B, C, Fig S3) for the genes expressed within these cell clusters). Particularly, an increased number of cells related to the cluster of inflamed epithelial cells was detected in CRC tissue compared to CRC adjacent intestinal tissue. This cluster of cells expressed a few epithelial cell markers, but additionally several neutrophil-associated markers such as S100A8 and S100A9, IL1B, MMP9, together with the neutrophil-recruiting chemokine CXCL8. Such clusters had described before in intestinal tissue^29^. In addition, CRC tissue samples exhibited a higher abundance of macrophages compared to their matched CRC adjacent intestinal tissues.

**Figure 2:**
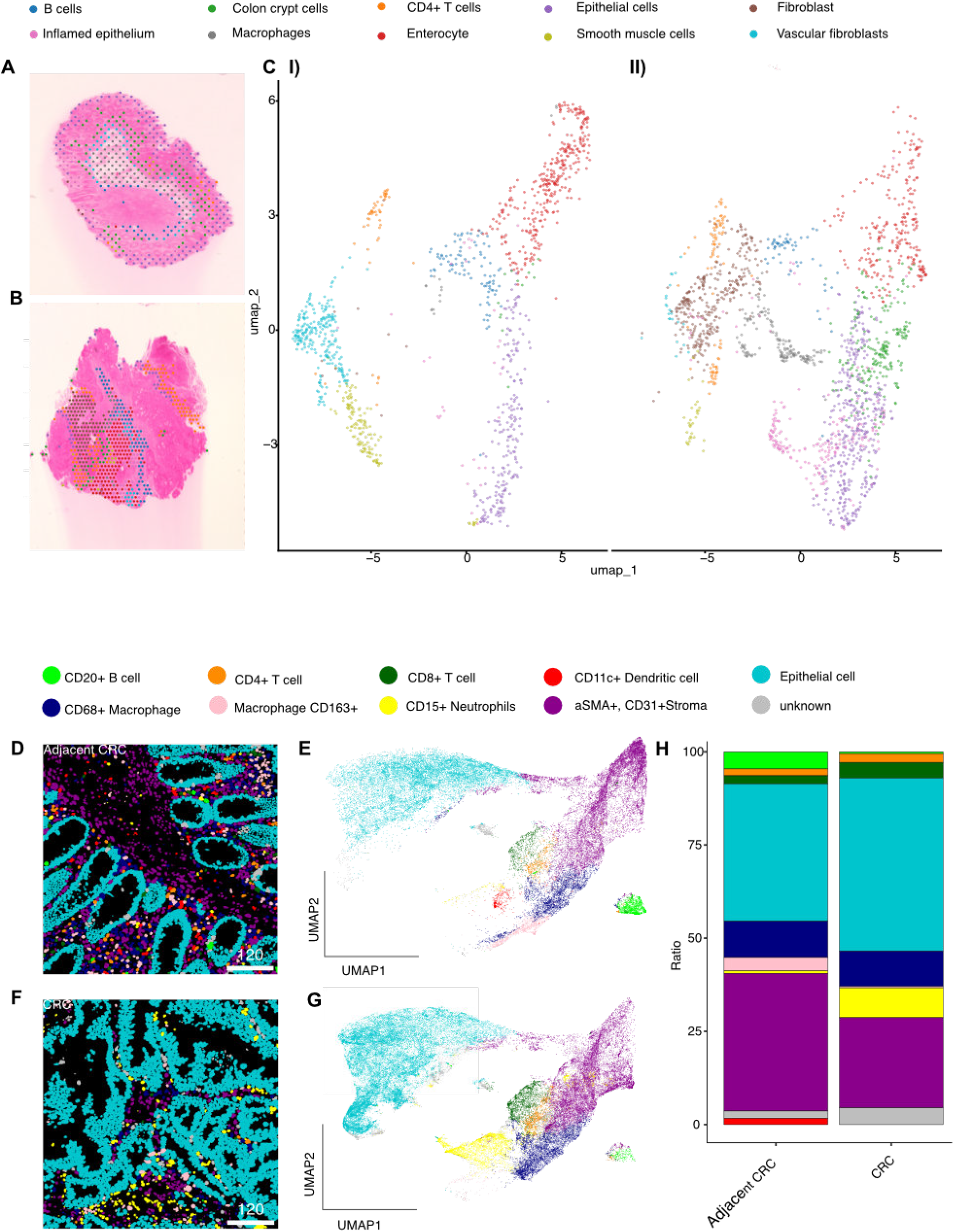
Immune cell types defined in 5 µm CRC adjacent intestinal tissue and CRC tissue. **(A):** Representative H&E staining and spatial annotation of 10 cell types detected in 3 CRC adjacent intestinal tissue and (**B)** matched CRC sections using spatial transcriptomics. **(C)** UMAP plot showing the distributions of 10 distinct cell types from (I) all CRC adjacent intestinal tissue and **(II)** CRC samples detected in spatial transcriptomics. **(D):** Representative imaging mass cytometry (IMC) image of 10 different cell types in CRC adjacent intestinal tissue, and (**E):** respective UMAP showing cell type clustering. (**F):** Representative IMC image of cell types in CRC and (**G):** Respective UMAP representing clustering of cell types. (**H)**: Ratio of cell types in CRC adjacent intestinal tissue and matched CRC tissue of 9 patients (two regions scanned per indication and patient n=18), quantified using IMC. Wilcoxon t-test was applied to determine differences in cell type composition between CRC adjacent intestinal tissue and CRC. Significant differences were considered at *p* < 0.05.

### Imaging mass cytometry detects a decrease of antigen-presenting cells accompanied by an increase in neutrophils in CRC tissue compared to adjacent tissue

To understand which immune cells are interacting with bacterial LPS in CRC adjacent intestinal tissue and in CRC tissue, we applied imaging mass cytometry (IMC) using a panel with 31 different immune cell markers as well as bacterial LPS (Suppl. Table S1). Two regions (each 600 µm x 600 µm) per CRC adjacent intestinal tissue (Fig. 2D, E) as well as CRC tissue (Fig. 2F, G) from nine patients including the patients with CRC studied above were analyzed. The mean cell count per image was lower in CRC adjacent intestinal tissue samples (2447 ± 321 SD) than in tumor samples (3140 ± 703 SD) (Fig. S4A). Overall, 44053 cells were identified in all adjacent tissues and 56522 cells in all CRC tissues (Fig. S4B). To define the cell types in adjacent tissues and CRC tissues, we applied an unsupervised clustering approach using FlowSOM, determining cell types based on similar marker expressions found within a cluster of cells (Suppl. Table S2, Figs. S5, S6, S7). The frequency of cell types from each of the two regions scanned per patient and tissue was comparable (Fig. S7). The most abundant cell populations were epithelial cells in both CRC adjacent intestinal tissues and CRC tissues, on average 36.8% and 46.8%, respectively (Fig. 2H, Fig. S8A). There were comparable frequencies of CD68+ macrophages (both 9%), CD4+ T cells (1.8 % and 2.2%), and CD8+ T cells (2.2% and 3.6%) in both CRC adjacent intestinal tissues and CRC tissues. Additionally, we detected a distinct CD68+ macrophage population, which was positive for CD163. CD163 is a plasma membrane glycoprotein and member of the scavenger receptor cysteine-rich (SRCR) superfamily class B, which is highly expressed in tissue-resident macrophages^30^. Here, the CD163+ macrophage population was significantly more abundant in CRC adjacent intestinal tissue (3.6%) compared to CRC tissue (0.42%). CD20+ B cells were decreased in CRC tissue (0.43%) compared to CRC adjacent intestinal tissue (4.58%) and, similarly, αSMA+, CD31+ stroma cells were found in a significantly lower ratio in CRC tissue (23.9%) than in CRC adjacent intestinal tissue (36.9%). CD11c+ dendritic cells (1.7%) were the least abundant cell type in CRC adjacent intestinal tissue. Of note, there were too few CD11c+ dendritic cells in the CRC tissue to form such a cluster cell type, resulting in the absence of this cell type in the CRC tissue based on these specific markers used in IMC. In contrast, CD15+ neutrophils were significantly increased in CRC (8.6%) compared to CRC adjacent intestinal tissue (0.64%). Interestingly, these neutrophils additionally expressed Granzyme B. The frequency of cell types from each of the two regions scanned per patient was comparable (Fig. S4). One tumor patient was almost purely positive for only two of the nine identified cytotoxic effector molecules, a type not typically associated with most neutrophil subtypes. Confirming our findings, our spatial transcriptomics and IMC data revealed overlapping cell populations, with B cells, epithelial cells, and inflamed epithelium showing consistent trends across both approaches, even though additional cell types were uniquely detected by each method (Fig. S8A and S8B).

### Distinct bacterial LPS-immune cell crosstalk in CRC adjacent intestinal tissue compared to CRC tissue

Having identified the cell types in the CRC adjacent intestinal and CRC tissues, we next investigated potential interactions between distinct human host cell types and bacterial LPS. We defined a cell as associated with bacterial LPS (LPS+) if the expression of the bacterial LPS marker within a cell exceeded the 90^th^ percentile cutoff in both the CRC adjacent intestinal tissue and the CRC tissue. Therefore, we chose a strict cutoff to minimize any potential background or overspill from different channels. Among the cells rated as associated with bacterial LPS, we observed a higher relative expression of activation markers, including CD25 in B cells, CD69 in CD4+ and CD8+ T cells, as well as CD163 in macrophages (Fig. S9).

By IMC, we observed that LPS signals appeared as more isolated dot-like spots in the CRC adjacent intestinal tissues, in contrast to a more diffuse signal in CRC tissues (Fig. 3A, B). However, we did not observe any statistical differences for the overall number of cells that were LPS+ between adjacent and CRC tissue (Fig. 3C). Subsequently, we investigated the relationship between the abundance of bacterial LPS+ cells and the ratio of distinct immune cell subsets (Fig. 3D). Linear regression analysis of tumor samples revealed a moderate positive relationship between the ratio of LPS+ cells and immune cells to all cells (regression coefficient β = 0.88, *p* = 0.06), indicating a trend towards increased immune cell infiltration with a higher overall count of LPS+ immune cells in the CRC tissue. In contrast, this association was not observed in the CRC adjacent intestinal tissue (β = 0.11, *p* = 0.74), highlighting a tumor-specific interaction. These findings suggest that higher proportions of LPS+ cells in CRC tumors may coincide with an increased immune cell infiltration into the CRC tissue. Finally, we assessed how frequently individual cell types were associated with bacterial LPS. Within the CRC adjacent intestinal tissue, the majority of dendritic cells were associated with bacterial LPS (65 %) (Fig. 3E). Similarly, on average 30% of all neutrophils and 26% of all CD163+ macrophages in CRC adjacent intestinal tissue were LPS+. For these three cell types, Kruskal Wallis test showed significant differences compared to the other cell types. Other ratios of LPS+ cells per cell type were below 12% within the CRC adjacent intestinal tissue. For the CRC tissue, the highest ratio of LPS+ cells per cell type was 16% in CD163+ macrophages, 14% in CD15+ neutrophils, and 13% in epithelial cells (Fig. 3F). All other LPS+ ratios per cell types were below 12 % in the CRC tissue. Kruskal-Wallis analysis only showed significant differences between epithelial cells and CD4+ T cells and B cells within the CRC tissue.

**Figure 3:**
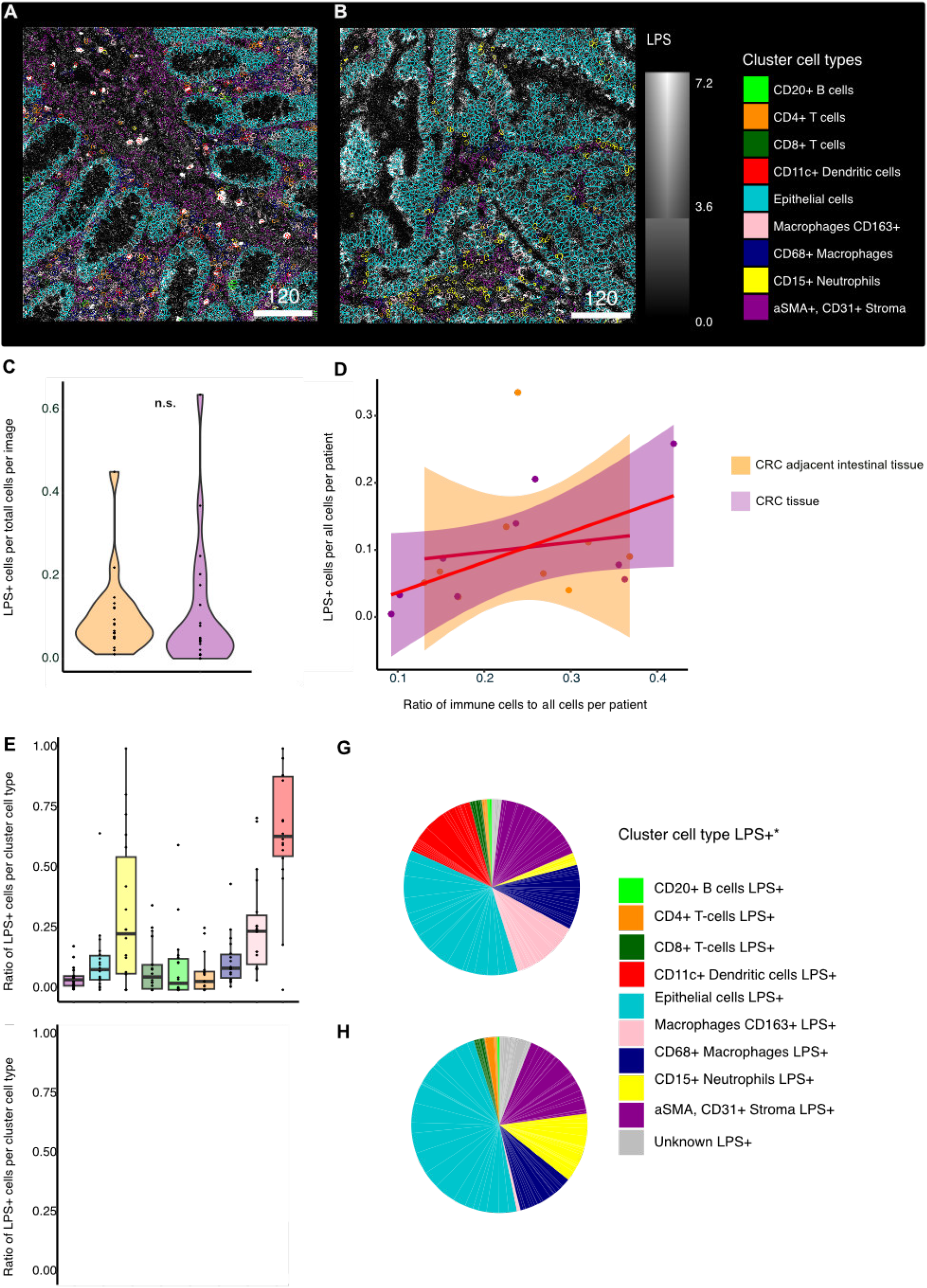
Differences in LPS-associated cell types between paired CRC adjacent intestinal tissue and CRC using IMC. **(A-B):** Representative images of LPS expression and associated cell types in CRC adjacent intestinal tissue (A) and CRC (B). (**C):** Violin plots showing the distribution of LPS^+^ cells in CRC adjacent intestinal tissue (orange) and matched CRC (purple) from 9 patients, n=18. **(D):** Linear regression model showing the relationship between immune cell infiltration and LPS^+^ cell ratio. Each data point represents the average of the two images analyzed per patient sample for paired CRC adjacent intestinal tissue and CRC tissue (n=9), respectively. **(E):** Ratio of LPS^+^ cells per cell type cluster in CRC adjacent intestinal tissue and **(F):** CRC tissue. Kruskal-Wallis test followed by Dunn’s post-hoc test with Benjamini-Hochberg correction was used to determine potential differences in the ratios of LPS+ cells among different cell types in CRC adjacent intestinal tissue and CRC. **(G):** A pie chart showing the ratio of each cell type within the LPS+ cell population in adjacent intestinal tissue, and **(H):** CRC tissue. Two images from n=9 paired CRC adjacent intestinal tissue and CRC tissues, respectively. Differences in the ratio of LPS^+^ cell types between CRC adjacent intestinal tissue and CRC tissue were tested using the Wilcoxon signed-rank test for each cell type, with Bonferroni correction for multiple comparisons. Significant differences were considered at *p* < 0.05. ns = not significant (*p* > 0.05).

Additionally, we investigated the cell type composition among all LPS+ cells in CRC adjacent intestinal tissue versus CRC tissue to determine which immune cell types were the most represented among them. Among all LPS+ cells per image, the fraction of dendritic cells and CD163+ macrophages was higher in CRC adjacent intestinal tissue (Fig. 3G) compared to CRC tissue (Fig. 3H). Additionally, a higher ratio of epithelial cells and neutrophils was detected among LPS+ cells per image in CRC compared to CRC adjacent intestinal tissue. CD163+ macrophages, CD15+ neutrophils, and CD11c+ dendritic cells differed significantly between LPS+ cells per image in the CRC compared to the CRC adjacent intestinal tissue.

Metagenomic analysis of snap-frozen CRC adjacent intestinal tissue and CRC tissue samples from the same patients showed no significant differences in the overall composition of bacterial genera (Fig. S10A)^18^. While we found an increased abundance of the CRC-associated *Fusobacterium nucleatum* in the CRC tissue, which is in line with several previous findings, this trend did not reach statistical significance in our study (Fig. S10B). Notably, *Lachnoclostridium* sp. YL32 was the only species that showed a statistically significant difference in relative abundance between CRC and CRC adjacent intestinal tissues (Fig. S10C). This finding is consistent with previous reports indicating an enrichment of *Lachnoclostridium* sp. YL32 in patients with colorectal adenomas compared to healthy controls^31^.

### A distinct spatial context of LPS+-associated immune cells differentiates CRC adjacent intestinal tissue from paired CRC

We observed differences in the ratios of specific cell types in CRC tissue compared to CRC adjacent intestinal tissue, and consequently, changes in bacterial LPS interactions with immune cells within CRC tissue. Next, we investigated the surrounding spatial context of cells associated with bacterial LPS in CRC adjacent intestinal and CRC tissues. All cell types showed self-interactions, suggesting spatial clustering. Epithelial cells were compartmentalized in CRC adjacent intestinal tissue and CRC tissue and were not found in spatial proximity to other cell types. In the CRC adjacent intestinal tissue, CD15+ neutrophils were found in spatial proximity to LPS+ CD4+ T cells, LPS+ CD11c+ dendritic cells, and CD163+ macrophages (Fig. 4A). In the CRC tissue, CD15+ neutrophils were additionally located in close proximity to LPS+ CD8+ T cells and CD68+ macrophages. CD163+ macrophages were found in spatial proximity to CD4+ T and CD8+ T cells as well as to neutrophils and other CD68+ macrophages in CRC adjacent intestinal tissue. In contrast, there was a lack of spatial proximity between CD163+ macrophages and CD4+ and CD8+ T cells in CRC tissue (Fig. 4B). However, the cellular proximity between CD15+ neutrophils and CD8+ and CD4+ T cells was reduced in CRC adjacent intestinal tissue (Fig.4C, D) compared to CRC tissue (Fig. 4C, E).

**Figure 4:**
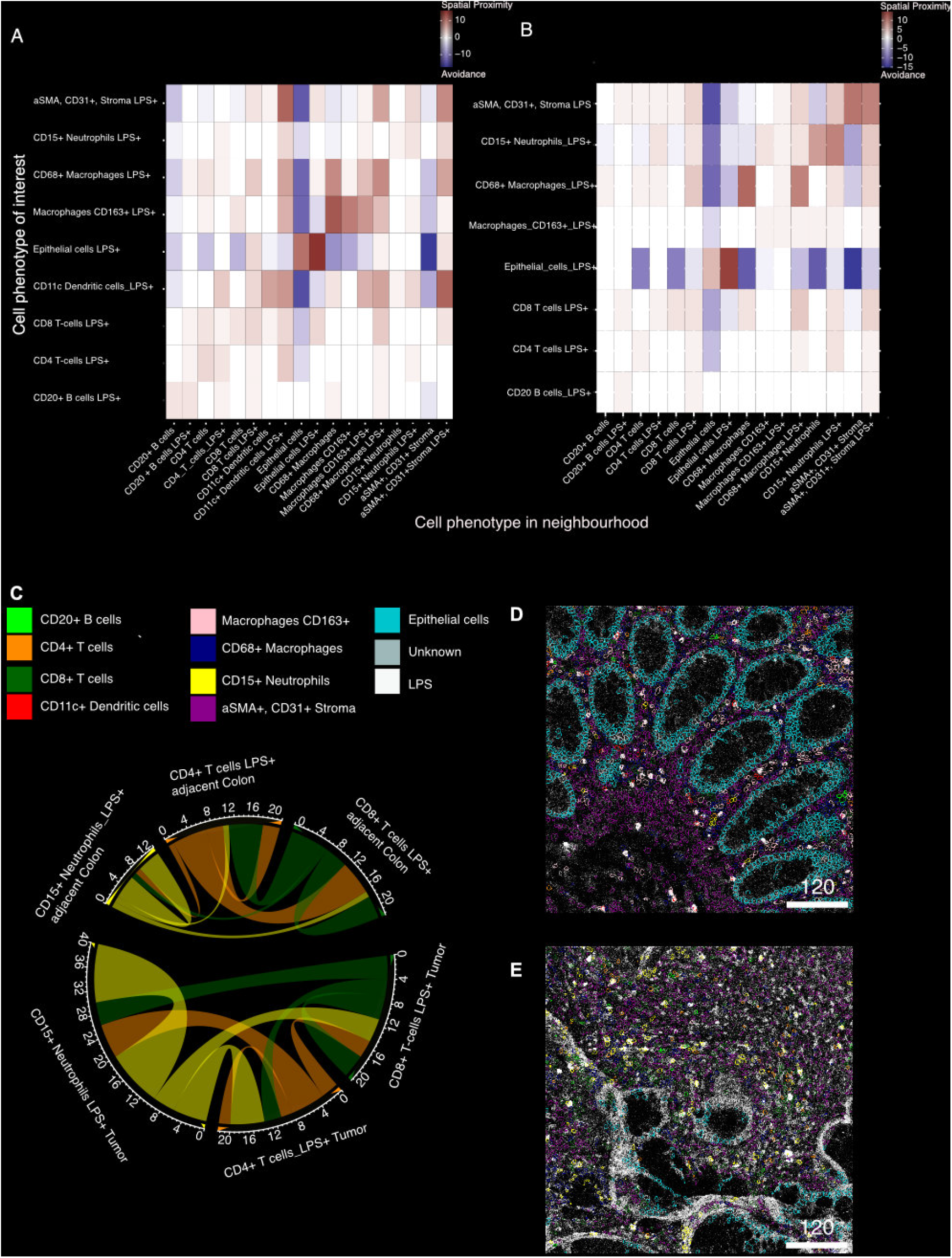
Differences in the spatial context of LPS-associated cells in CRC adjacent intestinal tissue and CRC tissue via IMC. **(A-B)**: Heatmap of spatial proximity observed in 9 (A) CRC adjacent intestinal tissue (B) and 9 CRC tissues, 2 images analyzed per individual patient tissue. Each cell type in each row is significantly neighbored (red) or avoided (blue) by the cell type in each column. Significance was determined using permutation test (P < 0.01). **(C):** Chord diagram summarizing cell-cell interactions of CD15+ neutrophils and CD4+ and CD8+ T-cells in CRC adjacent intestinal tissue and CRC tissue. **(D):** Exemplary IMC images showing CD15 neutrophil, CD8 and CD4 T cells interactions in CRC adjacent intestinal tissue, and **(E):** CRC tissue.

Overall, here we observed alterations in the composition of LPS–immune cells and in the spatial context, whereas no significant differences in bacterial profiles were detected between paired CRC and CRC adjacent intestinal tissues.

## Discussion

Our study provides evidence for shifts in the location of bacteria-derived LPS and bacteria-immune cell interactions from the CRC adjacent intestinal tissue to the paired CRC TME. Using 3D histology of cleared ∼5 mm^3^ tissues, we showed that bacteria-derived LPS can be found translocated to the subepithelial/submucosal tissue, both in the tumor as well as in the CRC adjacent intestinal tissue. Regional abundance of bacterial LPS was characteristic of CRC tissue. Additionally, we observed an overall higher bacterial biomass per mm^3^ in the CRC tissue compared to the CRC adjacent intestinal tissue. This finding is well in line with recent data^27,32^. However, we did not detect a significantly higher amount of bacterial LPS in the IMC data set, highlighting the heterogeneous distribution of bacteria-derived LPS in tumors. The localized patches of bacteria, as seen in our 3D imaging data, emphasize the importance of whole biopsy/tissue resection analysis for microbiome studies. Thus, standard 2D histology of 5 µm-thick FFPE sections might miss bacterial presence in tumor samples unless a multitude of serial sections are analyzed.

In the 3D histological data, we observed a high level of colocalization of bacterial LPS with immune cells. The investigation of immune cell types via spatial transcriptomics and IMC showed alterations in immune cell numbers in CRC tissue, along with an association of immune cell with bacterial LPS. First, CD11c^+^ dendritic cells were frequently associated with LPS in CRC adjacent intestinal tissue; however, this specific dendritic cell subtype was not detected within tumor samples, which might be due to the impact of tumor-secreting factors on the development and numbers of this dendritic cell subtype within tumors^18^. Furthermore, we observed a lower amount of CD163+ macrophages in tumor tissue compared to matched adjacent colon tissue, which is in line with findings in the literature^22^. Interestingly, CD163+ expressing macrophages are found in higher numbers in inflamed tissue; for example, over-expression of CD163 was found in areas of active inflammation in IBD, and the expression of both CD163 RNA and protein was increased in IBD in comparison to normal controls^23^. Here, we found that in adjacent colon samples, CD163+ macrophages were frequently associated with bacterial LPS. In line with this LPS+ association, CD163 can function as a macrophage receptor for bacteria, acting as an inducer of local inflammation during bacterial infection^24^.

Next, we observed CD15+ neutrophils already at the CRC adjacent intestinal site, a cell type not prevalently found in colon tissue under homeostatic conditions^33^. In fact, crosstalk with the commensal gut microbiota has been found to suppress neutrophil recruitment during intestinal health^34^. The presence of neutrophils at the CRC adjacent intestinal tissue could reflect an impaired gut barrier integrity^35^. A weakened gut barrier might subsequently enable bacterial translocation to the underlying layers, as supported by our 3D histology data of CRC adjacent intestinal tissue. Additionally, as seen in our study and supported by findings in the literature, neutrophils have been increasingly found within the CRC tissue^32^. Interestingly, the neutrophil population identified in our IMC data set co-expresses granzyme B, which is consistent with a previous study where this cell type was detected in human colon tumors and rodent models^36^. Lipid A analog, the lipid part of LPS, induced the release of granzyme B by these neutrophils, leading to tumor cell apoptosis^36^. Hence, the CD15+ granzyme B-expressing cell subtype population might represent an anti-tumor population of neutrophils, suggesting a pro-inflammatory N1-like type^36^. In line with this, our spatial transcriptomics data revealed that pro-inflammatory, activated neutrophil-associated genes were upregulated explicitly in CRC tissue. In contrast, tumor-associated neutrophils might also induce the shielding of cytotoxic immune cells from tumors due to their neutrophil extracellular trap (NET) formation^20,37^. In a recent study, immature CD66b+ cells, which are also used as a neutrophil marker, were found in immunosuppressive regions of CRC^15^.

Consistent with our findings, an increased infiltration of neutrophils and CD4^+^ T cells in CRC tissue compared to healthy mucosa was moderately associated with the presence of specific bacterial strains, such as *Bacteroides fragilis* and *Fusobacterium nucleatum*, and more strongly linked to a general increase in overall bacterial activity beyond these core pathogens^27^. Here, we see a higher association of CD15+ neutrophils with CD4+ T cells and CD8+ T cells in tumor tissue compared to the adjacent colon. Neutrophil interactions with T cells can lead to both suppression and apoptosis as well as activation^21,38-41^. Thus, in addition to spatial proximity, a more detailed analysis of the neutrophils and T cell markers could provide insights into their functional role and whether immunosuppressive or pro-inflammatory effects occur after interactions of neutrophils and T cells in CRC. Metagenomic analysis of paired CRC and adjacent tissue from the same patients revealed only one gram-positive bacterial species as differentially abundant. Thus, rather than individual bacterial species driving immune cell recruitment, the overall presence of translocated bacteria, bacteria-derived LPS, or other bacterial products may shape the immune landscape within the CRC tissue.

Overall, our findings contribute to the understanding of the heterogeneous TME and the bacteria as a component of it. We observed an inflammatory phenotype of the CRC tissue, accompanied by increased bacterial abundance in CRC tissue compared to paired adjacent non-tumor tissue. Moreover, distinct changes in specific immune cell populations and their associations with bacterial LPS were identified in CRC tumors. A deeper understanding of these bacteria–immune cell interactions will be essential for understanding CRC pathogenesis and finally advancing bacteria-based therapeutic strategies in CRC.

## Author contributions

Conceptualization: MS, BH

Data curation: ÅW, YM

Formal analysis: ÅW, YM

Funding acquisition: MS, AMR, BH

Investigation: ÅW, AMR, RZ, CM, CG, AW, MA, CB, MT, MR, PW, MW, LT, AA, AM, BH, YM, MS

Methodology: AMR, RZ, ÅW

Project administration: MS

Supervision: MS, BH

Visualization: ÅW

Writing – original draft: ÅW

Writing – review & editing: All authors

## Acknowledgments

Microscopy was performed with the equipment of the Center for Microscopy and Image Analysis. Spatial Transcriptomics was performed at the Functional Genomics Center Zurich (FGCZ) of the University of Zurich and ETH Zurich

AMR received funding from the Filling-the-Gap grant, University of Zurich. This work was supported by the Swiss Cancer League grant no. KFS-5372-08-2021-R (MS), a research grant from the Stiftung für wissenschaftliche Forschung an der Universität Zürich grant No STWF-22-009 (MS), Research grant from the ISREC Foundation (MS), Lighthouse project grant of the Comprehensive Cancer Center Zurich (CCCZ) und University Medicine Zurich (UMZH) (MS) and by the generous donors of the USZ Foundation (MS), Research grant from the Iten-Kohaut Foundation (BH), Research grant from the University of Zürich “Forschungskredit” (BH), Research grant from Novartis Foundation for medical-biological Research (BH), Research grant from the Fond’Action contre le cancer (BH).

## Declaration of Interest

MS has shares and is co-founder of Recolony/Adularia AG, Zurich, CH and has shares in PharmaBiome AG, Zurich, CH. MS served as Advisor for Abbvie, Gilead, Fresenius, Topadur, Takeda, Roche, Astra Zeneca, Pileje and Celltrion. MS received speaker’s honoraria from Janssen, Falk Pharma, Vifor Pharma, Pileje, Institut Allergosan and Bromatech. MS received research grants from Abbvie, Takeda, Gilead, Gnubiotics, Roche, Axalbion, Pharmabiome, Topadur, Basilea, MBiomics, Storm Therapeutics, LimmatTech, Zealand Pharma, NodThera, Calypso Biotech, Menarini, Pileje, Herbodee, Vifor. MT served as an advisor for Topadur and Takeda. MT received speaker’s honoraria from Janssen, Takeda and Intuitive Surgical. All other authors declare no competing interests relevant to this work.

## Resource availability

Raw TIFF image files generated from 3D histology and IMC experiments have been deposited at figshare https://doi.org/10.6084/m9.figshare.31054639 and are publicly available as of the date of publication. Metagenomic sequencing data from 9 CRC patients constitute a subset of a broader microbiome analysis previously published^18^ https://www.ncbi.nlm.nih.gov/bioproject/PRJNA1024674. Their respective Patient ID can be found in the supplementary table S3. This study did not generate new unique reagents or report original code. Requests for further information and resources should be directed to and will be fulfilled by the lead contact.

## Supplemental Material (separate file)

Supplementary Figure legends

Material and Methods

Supplementary Tables

## Notes

https://doi.org/10.6084/m9.figshare.31054639

