## Supplementary Material for "Altered Crosstalk of Bacterial Lipopolysaccharide with Immune Cells in Colorectal Cancer Compared to Paired Adjacent Intestinal Tissue"

### Supplementary Figures

**Supplementary Figure S1: Bacteria–immune cell interactions in CRC adjacent intestinal tissue visualized by 3D histology. (A):** Serial z-sections (1–4) from CRC adjacent intestinal tissue showing bacterial lipopolysaccharide (LPS) and CD45 and tissue autofluorescence (gray) highlighting epithelial crypt cells and subepithelial layers.

**Supplementary Figure 2: Immune cell and vasculature detection in CRC adjacent intestinal tissue and CRC tissue using 3D histology. (A):** Boxplot showing the percentage of vasculature (PVALP) colocalizing with lipopolysaccharide (LPS) in CRC adjacent intestinal tissue (orange) and CRC tissue (violet). Boxes represent the interquartile range, with the median indicated by a horizontal line. **(B):** Violin plot displaying the ratio of detected vasculature (PVALP) per $mm^3$ between CRC adjacent intestinal tissue (orange) and CRC tissue (violet). **(C):** Ratio of detected immune cells (CD45) per $mm^3$ between CRC adjacent intestinal tissue (orange) and CRC (violet). **(D):** Percentage of immune cells (CD45) colocalized with LPS. n= 6 CRC adjacent intestinal tissue and n=5 matched CRC samples. Lines connect values from CRC adjacent intestinal tissue and CRC tissue from the same patient.

**Supplementary Figure S3: Markers for Spatial Transcriptomics:** Dot plot showing the highly expressed markers for each cluster, where the color indicates the relative expression and the size of the dot indicates the proportion of expression inside each cluster.

**Supplementary Figure S4: Quantification of cell counts per image and total cell numbers per tissue type in IMC. (A):** Boxplots showing the individual cell counts per image obtained by IMC for each sample (n = 9 CRC adjacent intestinal tissue and matched CRC patient, 2 images scanned per patient). Boxes represent the interquartile range, with the median indicated by a horizontal line. **(B)**: Bar plot displaying the total cell count in CRC adjacent intestinal tissue (orange) and CRC tissue (violet).

**Supplementary Figure S5: Cell phenotyping in CRC adjacent intestinal tissue.** (**A**): Flow SOM clusters for CRC adjacent intestinal tissue samples, n=9. (**B**): Cell type cluster definition based on the marker expression within each cluster from (A). Cells were defined as follows: 1=aSMA+, CD31+ Stroma cells, 2=E-cadherin+ Epithelial cells, 3= CD20+B cells, 4=CD8+ T-cells, 5=CD15+Neutrophils, 6=CD4+ T-cells, 7=CD15+Neutrophils, 8=CD68+Macrophages, 9=unknown, 10=Cd11c+Dendritic cells, 11=CD8+T-cells, 12=unknown, 13=Macrophages CD163+, 14=CD20+B-cells, 15=CD20+B-cells, 16=unknown, 17=CD20+B-cells, 18=unknown. Unknown cell types represent cells with multiple overlapping marker expressions and no clear cell type distinguishable **(C):** Heatmap of marker expression in the cluster. Each column represents one cell. A fraction of the overall cell count was displayed in the heatmap (n= 7000 cells).

**Supplementary Figure S6: Cell phenotyping in CRC tissue.** (**A**): Flow SOM clusters for CRC samples, n=9. **(B):** Cell type cluster definition based on the marker expression within each cluster from (A). Cells were defined as follows: 1 = aSMA+, CD31+Stroma, 2 = E-cadherin+ Epithelial cells, 3 = Epithelial cells, 4 = CD15+ Neutrophils, 5 = aSMA+, CD31+Stroma, 6 = CD8+ T-cells, 7 = unknown, 8 = CD15+ Neutrophils, 9= CD20+B-cells, 10=CD8+ T-cells, 11 =unknown, 12=E-cadherin+Epithelial cells, 13=CD4+ T-cells, 14=CD68+Macrophages, 15=CD15+Neutrophils, 16= Macrophages CD163+, 17= unknown, 18=unknown. Unknown cell types represent cells with multiple overlapping marker expressions and no clear cell type distinguishable **(C):** Heatmap of marker expression in the clusters. Each column represents one cell. A fraction of the overall cell count was displayed in the heatmap (n=10000 cells).

**Supplementary Figure S7: Cell type ratios per image of each patient in IMC.** Percent of cells per cluster cell type across **(A):** CRC adjacent intestinal tissue, n=9, and (**B):** CRC tissue, n=9. Two regions per patient sample were scanned

**Supplementary Figure S8: Cell type cluster detected in IMC and spatial transcriptomics.** **(A):** Boxplots representing the Ratio of cell types per image, quantified in IMC in CRC adjacent intestinal tissue, n=9, and CRC tissue, n=9. Two regions per patient were scanned. Significant differences were considered at $p < 0.05$ and marked with *. **(B):** Boxplots representing the number of cells within each cluster in each condition for each sample, n=3 paired CRC adjacent intestinal tissue and CRC samples, quantified via spatial transcriptomics. Boxes represent the interquartile range, with the median indicated by a horizontal line.

**Supplementary Figure S9: Marker expression per cell type in IMC.** Dot plots represent the relative expression (z-score scaled, blue to yellow) and the percentage of cells expressing each marker gene in different cell types from **(A)**: CRC adjacent intestinal tissue and **(B)**: CRC tissue. Marker values were filtered such that only values above the 90th percentile were retained.

**Supplementary Figure S10: Bacterial profiles of snap-frozen CRC adjacent intestinal tissue and CRC tissue. (A)**: Relative abundance of bacterial species detected in snap-frozen CRC adjacent intestinal tissue and primary CRC tissue from the same 9 patients, analyzed by metagenomic sequencing. These 9 patients represent the same CRC patients as analyzed with IMC. **(B):** Relative abundance of *Fusobacterium nucleatum* in CRC adjacent intestinal tissue and CRC tissue**. (C):** Relative Abundance of Lachnoclostridium sp. YL32 in CRC adjacent intestinal tissue and the CRC tissue. Statistical differences were assessed with a paired Wilcoxon test, and P values were adjusted using FDR.

**Supplementary Video 1: 3D Visualization of bacteria-immune cell interactions in CRC adjacent intestinal tissue using 3D histology.** Video displaying the patient-derived CRC adjacent intestinal tissue stained for the immune cells (CD45), vasculature (PVALP), and bacterial lipopolysaccharide (LPS) as shown in Figure 1A.

**Supplementary Video 2: 3D Visualization of bacteria-immune cell interactions in CRC tissue using 3D histology.** Video displaying the patient-derived CRC tissue stained for the immune cells (CD45), vasculature (PVALP), and bacterial lipopolysaccharide (LPS) as shown in Figure 1B.

### Material and Methods

#### Patient material and ethical statement

Ethical approval for collecting patient material was received by the Cantonal Ethics Committee of the Canton Zürich (EK-1755/PB_2019-00169, BASEC-No.: 2019-02277). All Material was collected after prior written consent from all patients. Patients with CRC, diagnosed with Union for International Cancer Control (UICC) stages I-IV and a minimum of 18 years of age, were recruited from University Hospital Zurich (USZ). Surgical resections of CRC patients were assessed by a pathologist, and CRC adjacent intestinal tissue and CRC were identified and collected. The CRC adjacent intestinal tissue and CRC tissue were processed similarly as described in the following sections.

#### 3D histology

For tissue clearing and immunolabeling of ~ 5 $mm^3$ thick tumor CRC adjacent intestinal tissue, and CRC samples, an organic solvent-based DISCO protocol based on iDISCO[1] and aDISCO[2] was used. Fresh human CRC surgery samples (CRC adjacent intestinal tissue and CRC) were fixed in 4 % formalin overnight at 4°C, shaking. The next day, tissues were washed for several hours in PBS at room temperature (RT) until stored in 0.1 % sodium azide (NaN3) in PBS at 4°C. Tissues were dehydrated in ascending methanol (MeOH; 20%,40%, 60%, 80%) in ddH20 each for one hour, followed by 2 times 100% MeOH each for 1 hour at RT, and 1 hour at 4°C, shaking. Bleaching was performed in 10% hydrogen peroxide in MeOH overnight at 4°C, shaking. Samples were rehydrated in serial incubations of 80%, 60%, 40%, and 20% MeOH in ddH2O, followed by PBS, each for 1 hour at RT, shaking. Samples were permeabilized using 0.2% TritonX-100 in PBS 2 times each for 1 hour at RT, shaking, and 0.2% TritonX-100 + 10% dimethyl sulfoxide (DMSO) + 2.3% glycine + 0.1% NaN3 in PBS overnight at 37°C, shaking. Blocking was performed in 0.2% Tween-20 + 0.1% heparin (10 mg/ml) + 5% DMSO + 6% donkey serum in PBS for 1 day at 37°C. Samples were stained gradually with primary antibodies using goat-anti-bacterial LPS (PA1-73178), mouse-anti-human CD45 (CLO159), and rabbit-anti-human PVALP (83483), and secondary antibodies using donkey-anti-goat conjugated to AlexaFluor647 (Jackson, 705-605-003), donkey-anti-mouse conjugated to AlexaFluor594 (715-585-150), and donkey-anti-rabbit conjugated to AlexaFluor488 (711-545-152) in 0.2% Tween-20 + 0.1% heparin + 5% DMSO in PBS. All antibodies were applied three times per sample every third day at 37°C, shaking, to achieve a final dilution of 1:400 for LPS staining, 1:600 for CD45 staining, and 1:1000 for PVALP staining in 0.2% Tween-20 + 0.1% heparin + 5% DMSO + 0.1% NaN3 in PBS. After primary and secondary antibody staining, samples were washed each four times each in 0.2% Tween-20 + 0.1% heparin + 5% DMSO in PBS for at least 2 hours at RT, shaking. Then, samples were dehydrated in serial incubations of 20%, 40%, 60%, and 80% MeOH in ddH2O, followed by two times 100% MeOH, each for 1 hour at RT, shaking. Clearing was performed in 33% MeOH in dichloromethane (DCM) for 2.5 hours for CRC and 3 hours for CRC adjacent intestinal tissue followed by two times in 100% DCM each for 5 minutes for both CRC adjacent intestinal tissue and CRC at RT, shaking. Refractive index (RI=1.56) matching was performed in dibenzylether (DBE) at RT overnight, non-shaking. Afterwards, samples were stored at 4°C, covered from light.

**Light-sheet fluorescence microscopy**

Cleared samples were clamped into an adjustable custom-3D-printed holder and dipped into an immersion chamber containing DBE (RI=1.56). Imaging was performed using mesoscale selective plane illumination microscopy (mesoSPIM) V6 “Revolver” [3]in the Center for Microscopy and Image Analysis (ZMB) at the University of Zurich, Switzerland. Consistent across all imaging settings, an iChrome MLE laser system with 488 nm, 561 nm, and 647 nm excitation wavelengths was used in combination with a 405/488/561/640 quadruple-bandpass filter. For visualization of the tissue background for CRC adjacent intestinal tissue sample USZ_Pat_36, as shown in figure S1, we recorded the unstained channel with the 405 laser and the 405/488/561/640 quadruple-bandpass filter. Similar laser intensities were applied across all samples. Images were acquired using a Mitutoyo M Plan Apo 5×/0.14 objective (pixel size: 1.1 µm/pixel). The exposure time on the Hamamatsu ORCA camera was set to 20 ms.

**3D image visualization and analysis**

3D image stacks were saved in the tiff file format and then converted to ims file format using Imaris file converter (version 10.2.0). For visualization and analysis, Imaris (version 10.2.0) was used. To detect bacterial LPS protein, we used the ‘surfaces object’ function and operated the following algorithms: (1) apply ‘Background Subtraction (Local Contrast)’ mode and set ‘Diameter of Largest Sphere’ to 5 µm, (2) uniformly used an intensity threshold value of 2000 to segment bacterial LPS protein, and (3) set the ‘Minimal Number of Voxels’ to 10 for filtering-out noise. CD45+ cells were detected using the ‘surfaces object’ function with the following algorithms: (1) apply ‘Background Subtraction (Local Contrast)’ mode and set ‘Diameter of Largest Sphere’ as 5 µm, (2) uniformly use an intensity threshold value of 222 to segment CD45 cells, (3) enable intensity-based 'watershed object separation' with a diameter of 5 µm, and (4) set the 'minimal number of voxels' to 10. To assess colocalization of bacterial LPS with immune cells, we quantified the number of surfaces classified as LPS that had a distance below 1 $\mu m^2$ to surfaces classified as CD45. For vasculature detection based on PVALP staining, we used (1) a Gaussian smoothing filter with a surface detail of 1 µm, (2) an intensity-based threshold of 915, and (3) a 'minimal number of voxels' threshold of 10. To assess colocalization of bacterial LPS with the vasculature, we quantified the number of surfaces classified as LPS that had a minimal distance below 1 $\mu m^2$ to surfaces classified as vasculature. To estimate the volume of each tissue sample, we used the autofluorescence signal in the 488 nm channel to distinguish the tissue from the background. Surface reconstruction was performed in Imaris using the surface tool, with a Gaussian smoothing level of 10 µm (surface detail parameter). This high degree of smoothing intentionally suppressed fine morphological features, yielding a simplified surface representation for approximating total tissue volume. Finally, the animation and snapshot functions in Imaris were used to export images and supplementary videos.

**Statistics 3D histology**

We performed a Wilcoxon rank-sum test to compare the measurements between CRC adjacent intestinal tissue and CRC tissue. The analysis revealed no statistically significant difference between the two groups.

**Validation of LPS and CD45 antibody specificity via 2D histology**

The specificity of the LPS and C45 antibody used in 3D histology was further evaluated on 5 µm FFPE tissue sections. Representative images from this validation are shown in Supplementary Figure S9. Paraffin-embedded tissue sections were subjected to deparaffinization and antigen retrieval prior to immunostaining. Briefly, slides were washed twice in Histo-clear (2 × 10 min) followed by graded ethanol washes (2 × 1.5 min in 100%, 2 × 1 min in 96%, and 1 × 2 min in 70%). Slides were then rinsed in Milli-Q water for 30 s. Antigen retrieval was performed by incubating slides in citrate buffer (pH 6.0) at 97 °C for 25 min, followed by gradual cooling to RT for 15–20 min. Two washes in PBS (2 × 10 min) and permeabilization in 0.1% Triton X-100 (10 min) followed. Blocking was performed with 0.2% Tween-20, 0.1% heparin, 5% donkey serum, and 6% DMSO in PBS for 2 h at RT. Primary antibodies, goat-anti-bacterial LPS (PA1-73178) and mouse-anti-human CD45 (CLO159) were each applied 1:400 in in iDISCO staining buffer (0.2% Tween-20 + 0.1% heparin + 5% DMSO in PBS) and incubated overnight at 4°C. On the following day, slides were washed with 0.2% Tween-20/0.1% heparin/PBS (2 × 20 min), then incubated with secondary antibodies, donkey-anti-goat conjugated to AlexaFluor647 (Jackson, 705-605-003) and donkey-anti-mouse conjugated to AlexaFluor594 (715-585-150), diluted 1:400 in 0.2% Tween-20 + 0.1% heparin + 5% DMSO in PBS for 2 h at room temperature. Subsequently, slides were washed in 0.2% Tween/0.1% heparine/PBS 1 × 20 min and 1 × 10 min. Then, nuclei were counterstained with DAPI (1:1000 in PBS, 5 min), followed by 5 min. wash in PBS and one in Milli-Q water. Slides were mounted with fluorescence mounting medium and allowed to set overnight at RT before storage at 4 °C until imaging.

Confocal laser scanning microscopy was performed on a Leica STELLARIS 5 inverted confocal microscope (DMi8 stand) operated with LAS X (version 5.0.3.24880, Leica Microsystems). Images were acquired using a 20×/0.75 DRY objective (HC PL APO CS2). Excitation was provided by the White Light Laser (WLL) and a 405-nm diode for DAPI. Emission was detected with Power HyD S detectors using the following detection windows: 420–595 nm (DAPI), 595-652 nm (AF 591) and 657-808 nm (AF 647). All datasets were exported as .lif and analyzed in ImageJ (v 1.54f) with identical processing across conditions.

**Spatial transcriptomics**

Spatial transcriptomics was performed using the Visium platform (10x Genomics) at the Functional Genomics Center Zurich. Raw data were processed in R (v.4.3) using the Seurat (v5.1) framework. Individual samples were imported as Seurat objects and merged into a single object. Quality control filtering was applied to exclude low-quality spots, removing spots with fewer than 200 or more than 10,000 detected genes (nFeature_Spatial), fewer than 800 or more than 60,000 unique molecular identifiers (nCount_Spatial), or with ≥20% mitochondrial transcript content (percent.mt). The filtered dataset was normalized using SCTransform, and dimensionality reduction was performed with principal component analysis (PCA) followed by Uniform Manifold Approximation and Projection (UMAP). Graph-based clustering was performed in Seurat, and clusters were visualized in UMAP space, stratified by sample and condition (Healthy vs. Tumor). Cluster annotation was guided by canonical marker expression and refined using SingleR with reference datasets from celldex. Annotation was further curated manually based on expression of key immune and stromal markers. Cluster identities were exported, and cell counts per cluster and condition were summarized by estimating the relative abundance for all the annotated clusters within one sample.

#### Imaging Mass Cytometry Staining on FFPE sections

FFPE sections were cut to 5 µm sections using a rotary microtome (Zeiss Hyram M 15) one day before staining and baked at 62°C for at least 2 h. For IMC staining, we followed the protocol Imaging Mass Cytometry Staining for FFPE sections from Fluidigm (PN400322 A3) according to the manufacturer's instructions. In brief, slides were dewaxed in xylene for 20 min, followed by descending grades of ethanol (100%, 95%, 80%, 70%) for 5 min each. Slides were washed in Maxpar water (201069 Fluidigm) for 5 minutes. Pre-warmed Antigen retrieval solution at pH 9 (S236784-2, Agilent) was applied at 96°C for 30 minutes. Afterwards, the slides were cooled to 70°C, followed by washing with Maxpar Water and Maxpar PBS (201058, Fluidigm) for 10 minutes each. Slides were blocked with 3% BSA (A3059, Sigma-Aldrich) in Maxpar PBS for 45 minutes at room temperature in a hydration chamber. Metal-tagged antibodies were applied to the slides (see supplementary table 1 for specific dilution of antibodies) and left overnight at 4°C in a hydration chamber. On the next day, slides were washed two times in 0.2 % Triton X-100 (85111, Thermo Scientific) in Maxpar PBS for 8 minutes. Cell-ID Intercalator-Ir (201192A, Fluidigm) in Maxbar PBS was applied for 30 minutes at room temperature, followed by a wash in Maxpar water for 5 minutes. Lastly, the slides were air dried for 20 minutes at room temperature. Stained slides were scanned using HyperIOn Imaging System. Two areas of 600 µM x 600 µM were scanned per patient sample.

#### IMC Data analysis

##### Image pre-processing (Single-cell quantification and identification of cell neighborhood)

Raw MDC files were processed using HistoCAT web version 2.1.4. Whole-cell segmentation was performed using Mesmer with E-cadherin as the cytoplasmic marker and DNA2 for the nuclear marker. Further processing was performed using the Steinbock docker framework, version 0.16.0[4]. In brief, images were processed to tiff format, and hot pixel filtering was applied to remove IMC-specific noise. Mean pixel intensities per cell and marker were extracted. Morphological properties were extracted per cell such as cell area, centroid, major and minor length of axes, and eccentricity. Cell neighbors were identified based on Euclidean distances between object borders.

To this end, pixel expansion was performed, and touching objects (i.e., objects within a 4-neighborhood) were considered as neighbors.

imcRtools version 1.8.0, was used to read in the Steinbock generated data into R, version 4.3.3 as a SpatialExperiment Object. Cytomapper version 1.14.0 [5] was applied to read in multi-channel images and segmentation masks in a cytoIMAGElist. Further IMC analysis was based on the workflow published by Windhager et al.[4]. Single-cell expression data were ashin transformed with a cofactor of 3 to avoid bias by highly expressing cells.

**Cell clustering in IMC**

The scater package, version 1.30.1 was used for a UMAP dimensionality reduction, and a seed was used to ensure reproducibility. The data was then visualized using patchwork, version 1.2.0 dittoSeq, version 1.14.0 and viridis packages, version 0.6.5. Single-cell marker expression was shown with a heatmap of a subset of 7000-10000 cells from the whole dataset using dittoHeatmap function (Fig. S5 and S6). As we first observed, clustering of cells based on the CRC patient IDs, we performed a batch correction based on mutual nearest neighbors (MNN) using the batchelor package, version 1.18.1. Nuclei and markers expressed in several cell types were excluded from clustering.

**Cell phenotyping in IMC**

The dataset was split into CRC adjacent intestinal tissue and CRC to determine cell clusters present in either. The CATALYST package, version 1.26.0, was used to perform self-organizing maps (SOM) clustering via ConsensusClusterPlus. The plot-delta_area function from the CATALYST package was applied to define the optimal number of clusters. Detected cluster cell types were manually annotated based on the marker expression within those clusters as observed in a heatmap and the single marker expression displayed in UMAP (Fig. Supplement 5 and S6). The main cell types featuring selective markers are displayed in table S2.

**Visualization of IMC images**

Multiplex imaging data were visualized using the cytomapper R package. Pixel-level intensities of the LPS marker were displayed with the plotPixels function, with similar brightness–contrast–gamma (BCG) adjustments applied to all images. The plotCells function was used to visualize segmentation-based single-cell data, with each cell colored according to its assigned cell type.

**Bacterial LPS analysis for IMC**

Phenotyped cells were assessed for their association with LPS marker expression. A cell was classified as $LPS^+$ if the expression of bacterial LPS exceeded the 90th percentile cutoff in CRC adjacent intestinal tissue and CRC, respectively. A dittoDotPlot from dittoSeq function in R was used to display the expression and the percent expression of all markers, exceeding the 90th percentile cutoff, among LPS positive and LPS-negative cell types (Supplementary Figure S9). Linear regression was used to investigate the relationship between immune cell infiltration and LPS-positive cell proportions per patient sample. The ratio of immune cells per patient sample was calculated by excluding epithelial cells, stroma cell and the unknown cell types from the analysis. Linear regression was applied using the lm() function in R, version 4.3.3. Model performance was evaluated by examining the regression coefficients and p-values obtained from the model summary.

**Spatial Proximity/ "Interactions" analysis used in IMC analysis**

Pairwise interaction between all cell types of the dataset was performed using the imcRtools package testInteractions function, which applies the interaction testing strategy by Schapiro et al. [6]

**Statistics applied in IMC analysis**

For analysis of cell type distribution between CRC adjacent intestinal tissue and CRC tissue and differential ratios of cell types among LPS+ cells in CRC adjacent intestinal tissue and CRC tissue, we used a Wilcoxon signed-rank test for each cell type separately, followed by multiple testing correction using Bonferroni. using the R stats package, 4.3.3. We applied a Kruskal-Walli's test followed by Dunn's post-hoc test with Benjamini-Hochberg correction using dunn.test package, version 1.3.6, to determine if there were significant differences in the proportion of LPS-positive cells per cell type for CRC adjacent intestinal tissue and CRC tissue, respectively. Lastly, a permutation test ($P < 0.01$) using the imcRtools package was applied with a BiocParallel package, version 1.36.0 to test for significantly neighboring (red) or avoided (blue) cell-cell proximity.

### Metagenomic Sequencing Analysis

Metagenomic sequencing of snap-frozen CRC adjacent intestinal tissue and CRC tissue from 9 CRC patients who were also subjected to IMC analysis was performed. These CRC samples represent a subset of a larger metagenomic dataset that has been published previously[7].The corresponding Patient IDs are provided in Supplementary Table S3.

### Supplementary Tables

#### Supplementary Table 1: Applied dilutions and metal tags of antibodies used for IMC.

| Target | Metal | Dilution factor | Clone | Company and Catalogue number |
|---|---|---|---|---|
| CD3 | 170Er | 100 | UCHT1 | Fluidigm, 3170019D |
| CD8a | 162Dy | 100 | D8A8Y | Fluidigm, 3162035D |
| CD31 | 151Eu | 200 | EPR3094 | Fluidigm, 3151025D |
| aSMA | 141Pr | 200 | 1A4 | Fluidigm, 3141017D |
| E-Cadherin | 158Gd | 100 | 24E10 | Fluidigm,3158029D |
| CD11c | 154Sm | 200 | Polyclonal | Fluidigm, 3154025D |
| CD45RO | 173Yb | 200 | UCHL1 | Fluidigm, 3173016D |
| CD45RA | 166Er | 200 | HI100 | Fluidigm, 3166031D |
| CD25 | 175Lu | 50 | EPR6452 | Fluidigm, 3175036D |
| HLA-DR (MHC class II) | 174Yb | 400 | LN3 | Fluidigm, 3174025D |
| CD194/CCR4 | 149Sm | 400 | L291H4 | Fluidigm, 3149030D |
| CD196 (CCR6) | 163Dy | 1000 | G034E3 | Fluidigm, 3163029D |
| Granzyme B | 167Er | 200 | EPR20129-217 | Fluidigm, 3167021D |
| CD86 | 165 Ho | 50 | IT2.2 | Fluidigm, 3176026D |
| CD68 | 159Tb | 100 | KP1 | Fluidigm, 3159035D |
| Ki-67 | 168Er | 100 | B56 | Fluidigm, 3168022D |
| CD4 | 156Gd | 50 | EPR6855 | Fluidigm, 3156033D |

| | | | | |
|---|---|---|---|---|
| CD16 | 146 Nd | 100 | EPR16784 | Fluidigm, 3146020D |
| CD49a/integrin alpha 1 | 172 Yb | 50 | A-9 | Santa Cruz, sc-271034 |
| CD69 | 142Nd | 50 | Polyclonal | Invitrogen, PA5-95721 |
| CTLA-4/CD152 | 176Yb | 100 | F-8 | Santa Cruz, sc-271034 |
| LPS | 153Eu | 50 | GNE11-265.5 | Abnova, MAB6173 |
| MBP | 152S m | 100 | MBP101 | Minvitrogen, A1-10837 |
| PDL1 | 150Nd | 50 | SP142 | Fluidigm, 3150033D |
| CD163 | 169T m | 100 | 5C6-FAT | Novus Biologicals, BM4041 |
| FOXP3 | 155Gd | 400 | 236A/E7 | Fluidigm, 3155016D |
| CD20 | 161Dy | 100 | H1 | Fluidigm, 3161029D |
| CLA | 143Nd | 100 | HECA-452 | Biolegend, B289471 |
| CD1a | 164Dy | 200 | CL703217 | Biotechne, MAB7076 |
| CD15 | 160Gd | 100 | W6D3 | Biolegend, B321296 |
| CD14 | 144Nd | 200 | EPR3653 | Fluidigm, 3144025D |
| CD206 | 171Yb | 200 | 15--2 | Biolegend, 321127 |
| ICSK1 | 195pt | 400 | IMC Cell Segmentation Kit | Fluidigm, TIS-00001 |
| ICSK2 | 196pt | 400 | IMC Cell Segmentation Kit | Fluidigm, TIS-00001 |

**Supplementary Table 2*:* Cluster cell type definition applied in IMC as seen in the heatmap in the supplementary figures S5 and S6.**

| **Cluster cell type** | **Marker Expression** |
|---|---|
| B-cell | CD20+, CD45RA/CD45RO |
| CD4+ T-cell | CD3+, CD4+, CD8- |
| CD8+ T-cell | CD3+, CD8+, CD4-, |
| Dendritic cell | CD11c+ |
| Epithelial cell | E-cadherin+, Ki67+ |
| Macrophage | CD68+, HLA-DR+, CD16+, CD14+ |
| Macrophage CD163+ | CD68+, CD163+, HLA-DR+ |
| Neutrophil | CD15, CD14+, Granzyme B+, CD68+ |

| | |
|---|---|
| Stroma | aSMA+, CD31+, E-cadherin- |

**Supplementary Table S3:** Patient ID of snap frozen CRC adjacent intestinal tissue and CRC tissue analyzed via metagenomic sequencing in Morsy et al. [7]

| Patient_ID | Material |
|---|---|
| USZ_Pat_24 | CRC adjacent intestinal tissue<br>CRC |
| USZ_Pat 25 | CRC adjacent intestinal tissue<br>CRC |
| USZ_Pat_33 | CRC adjacent intestinal tissue<br>CRC |
| USZ_Pat_34 | CRC adjacent intestinal tissue<br>CRC |
| USZ_Pat_35 | CRC adjacent intestinal tissue<br>CRC |
| USZ_Pat_36 | CRC adjacent intestinal tissue<br>CRC |
| USZ_Pat_37 | CRC adjacent intestinal tissue<br>CRC |
| USZ_Pat_54 | CRC adjacent intestinal tissue<br>CRC |
| USZ_Pat_131 | CRC adjacent intestinal tissue<br>CRC |
