## Supplementary figures and images for "Altered Crosstalk of Bacterial Lipopolysaccharide with Immune Cells in Colorectal Cancer Compared to Paired Adjacent Intestinal Tissue"

### Supplementary Figures

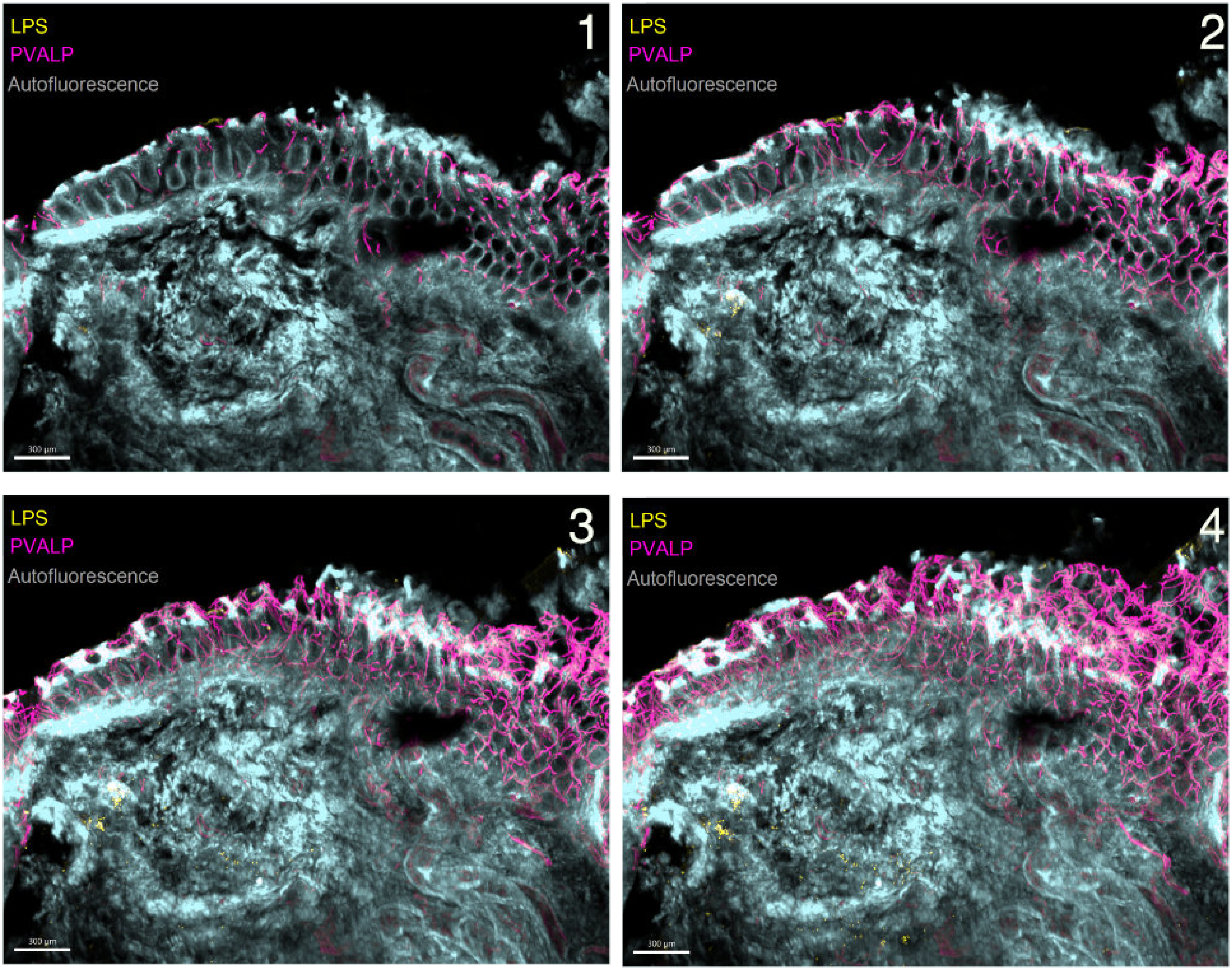

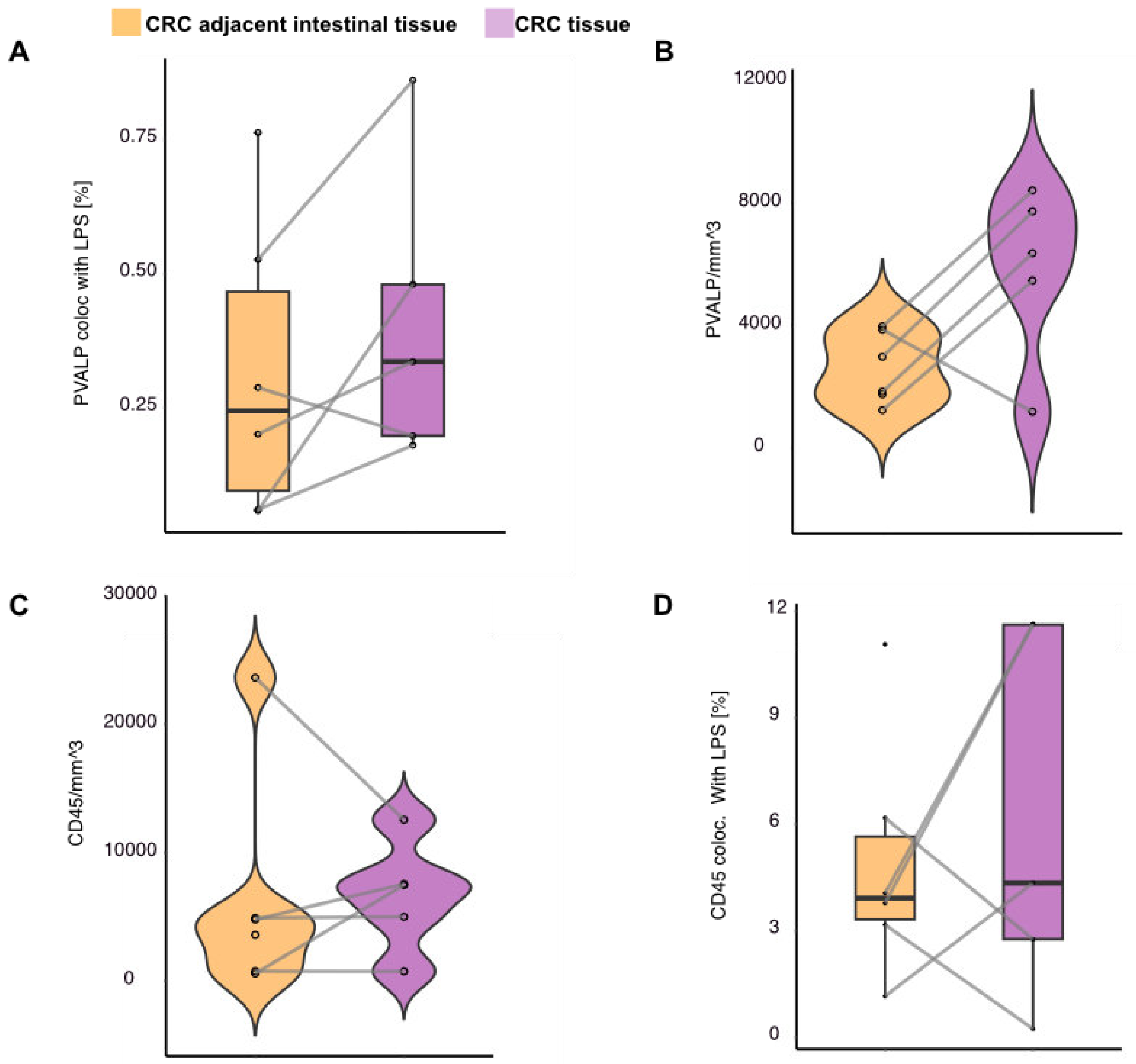

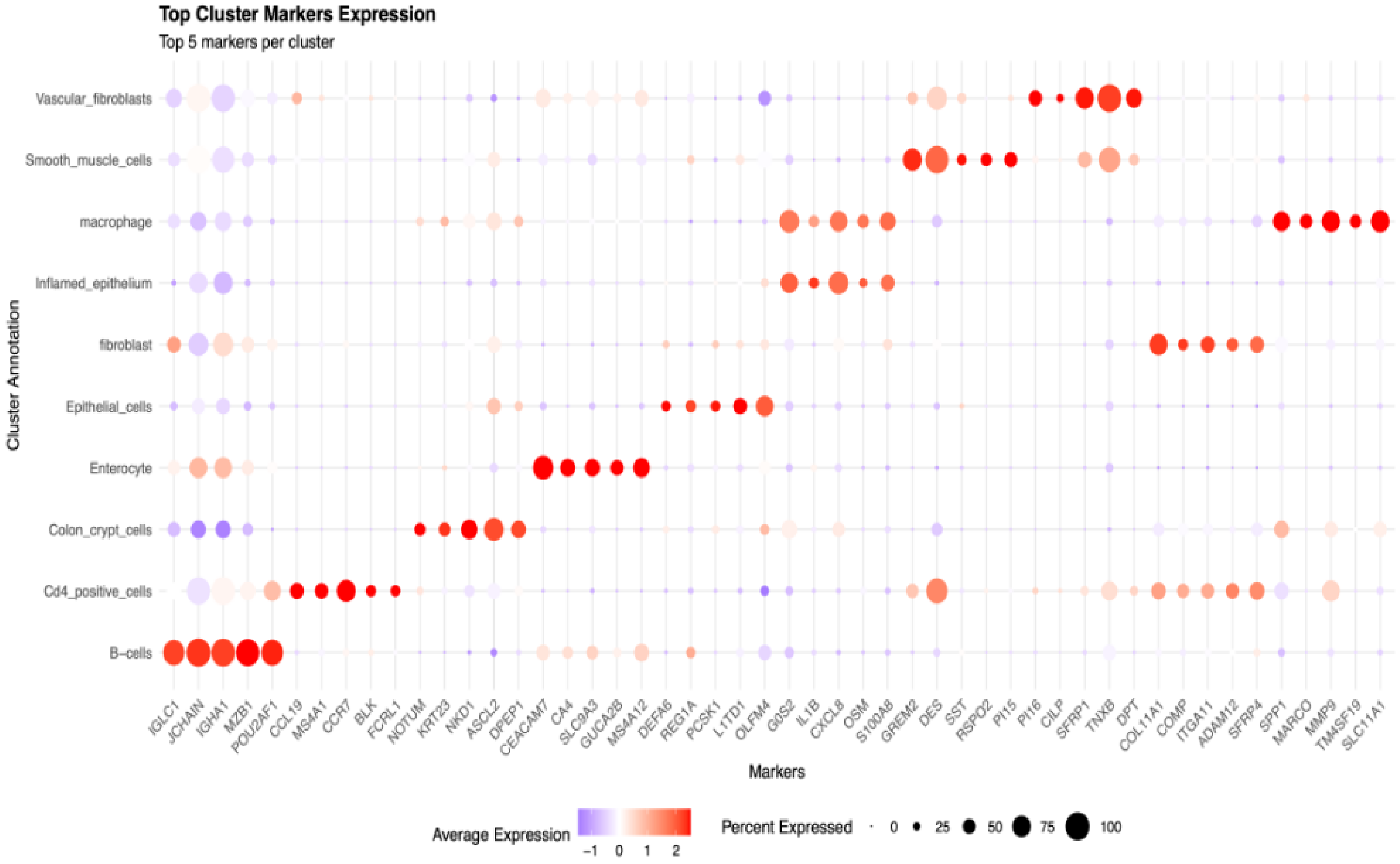

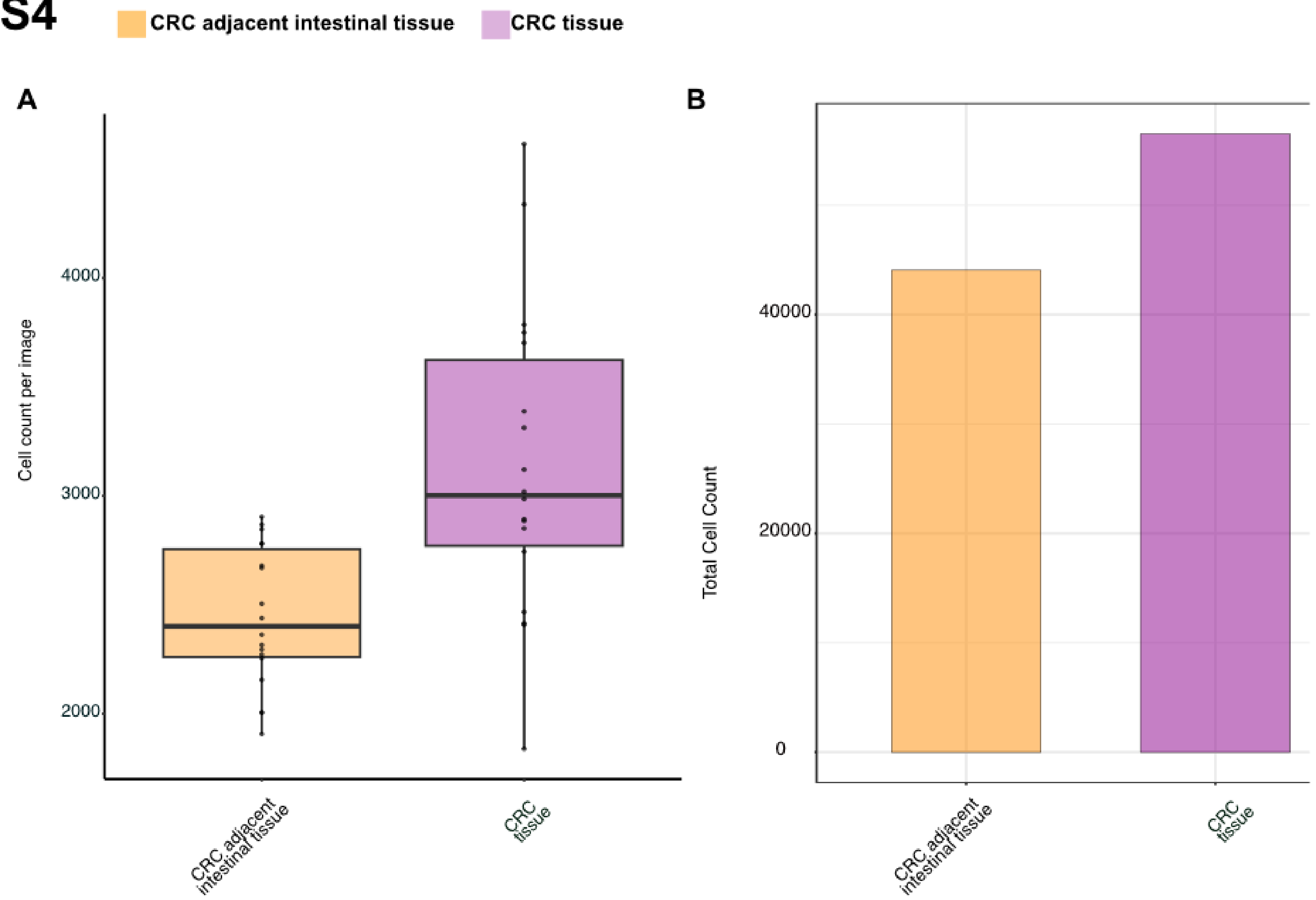

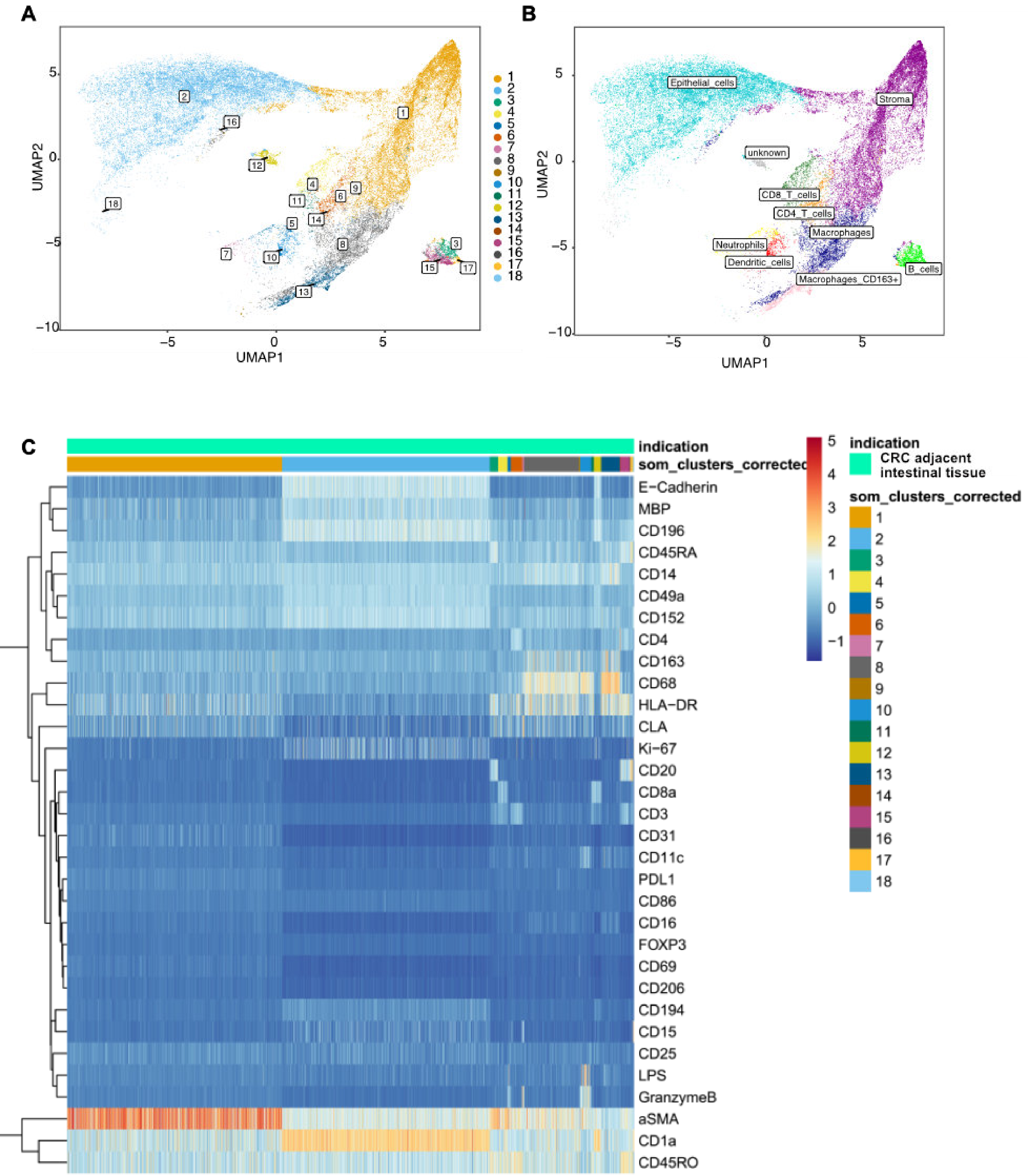

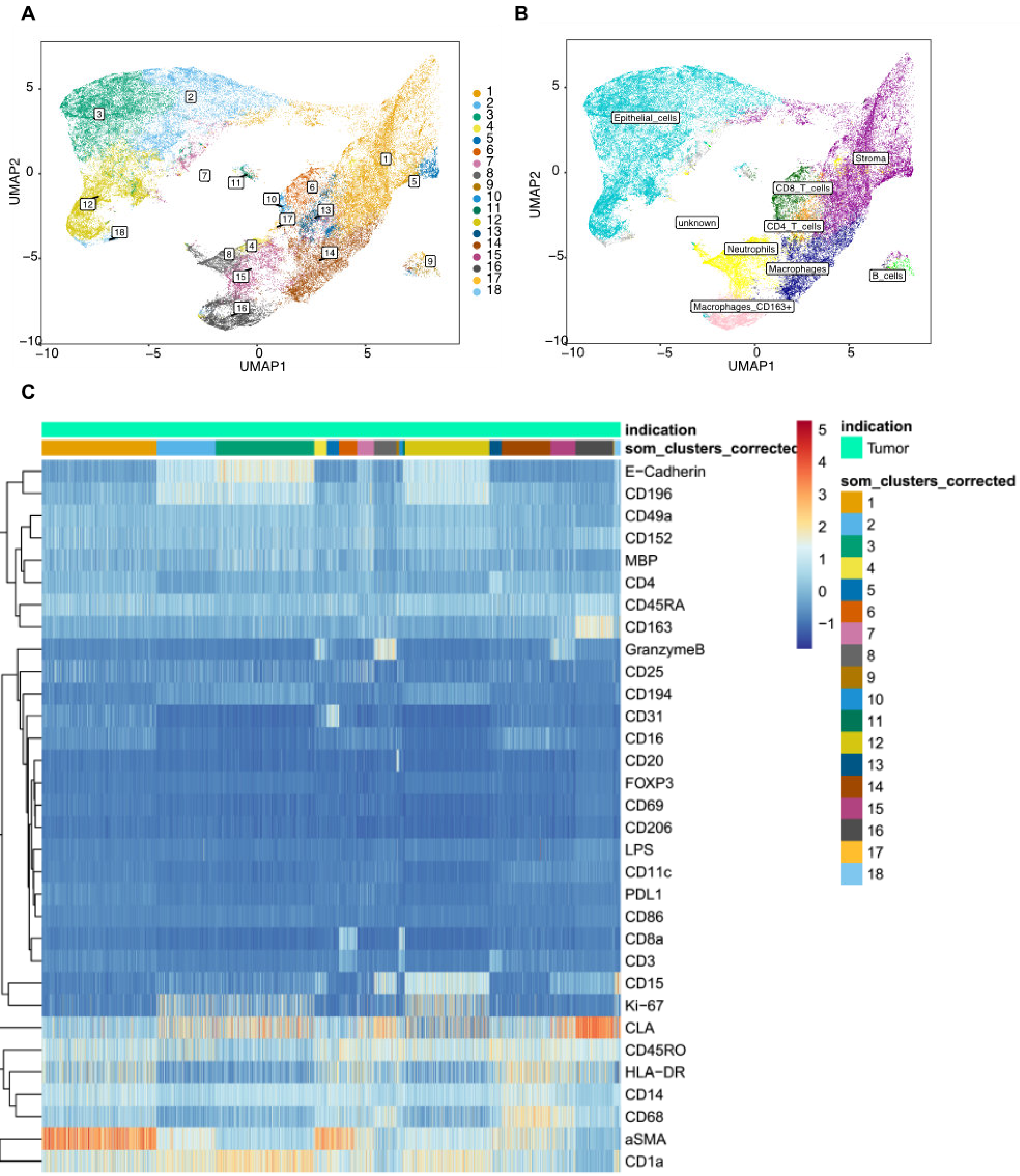

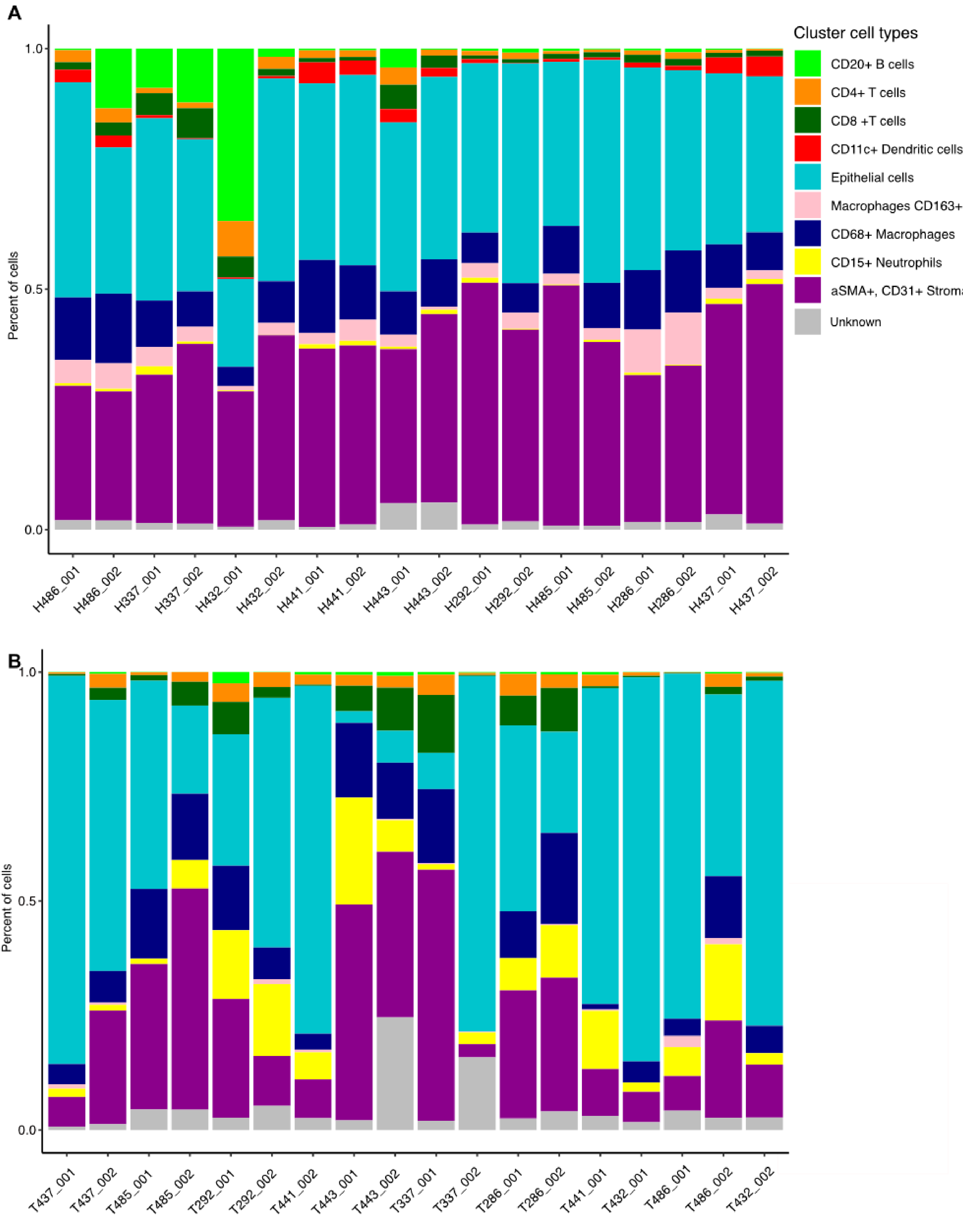

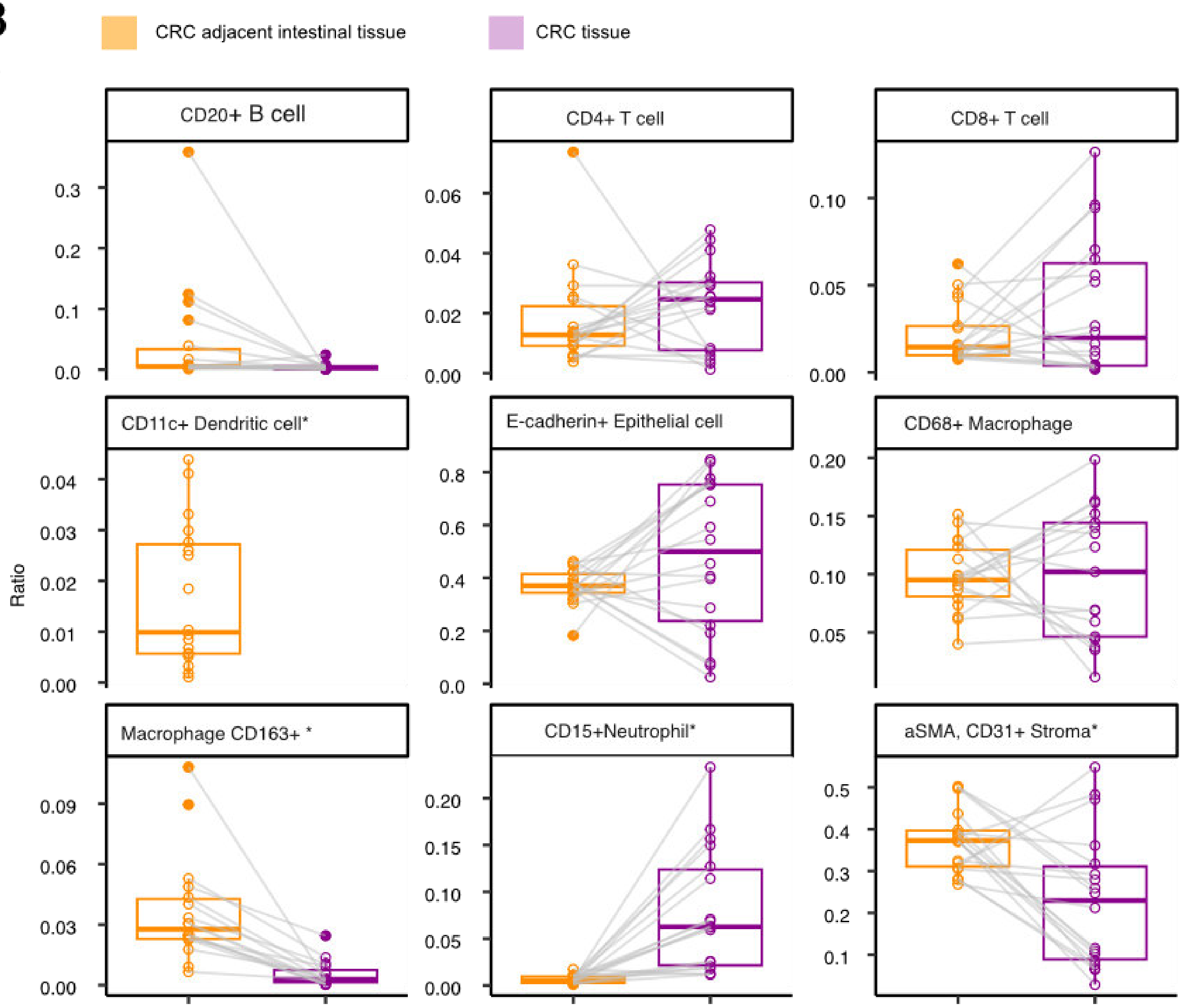

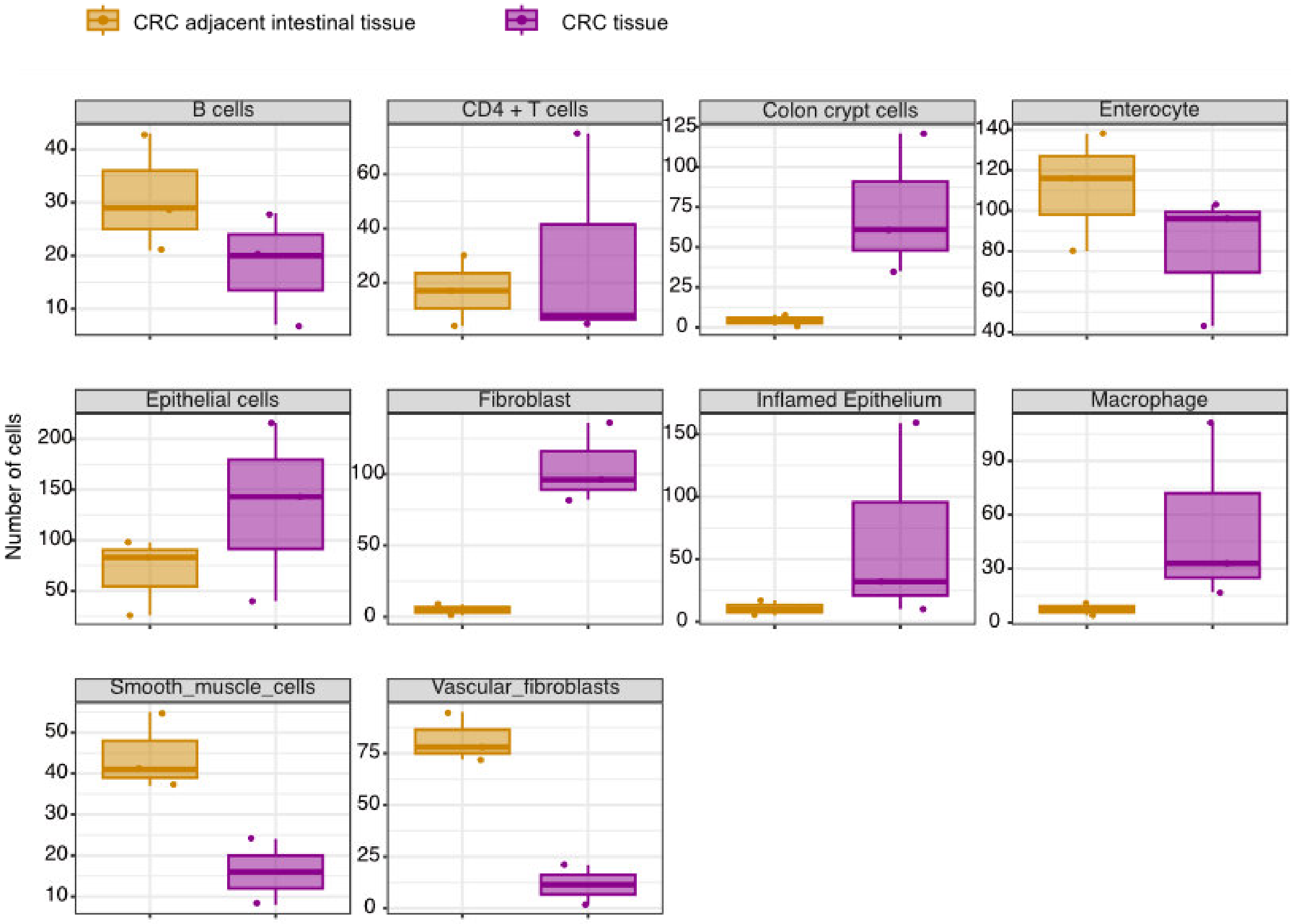

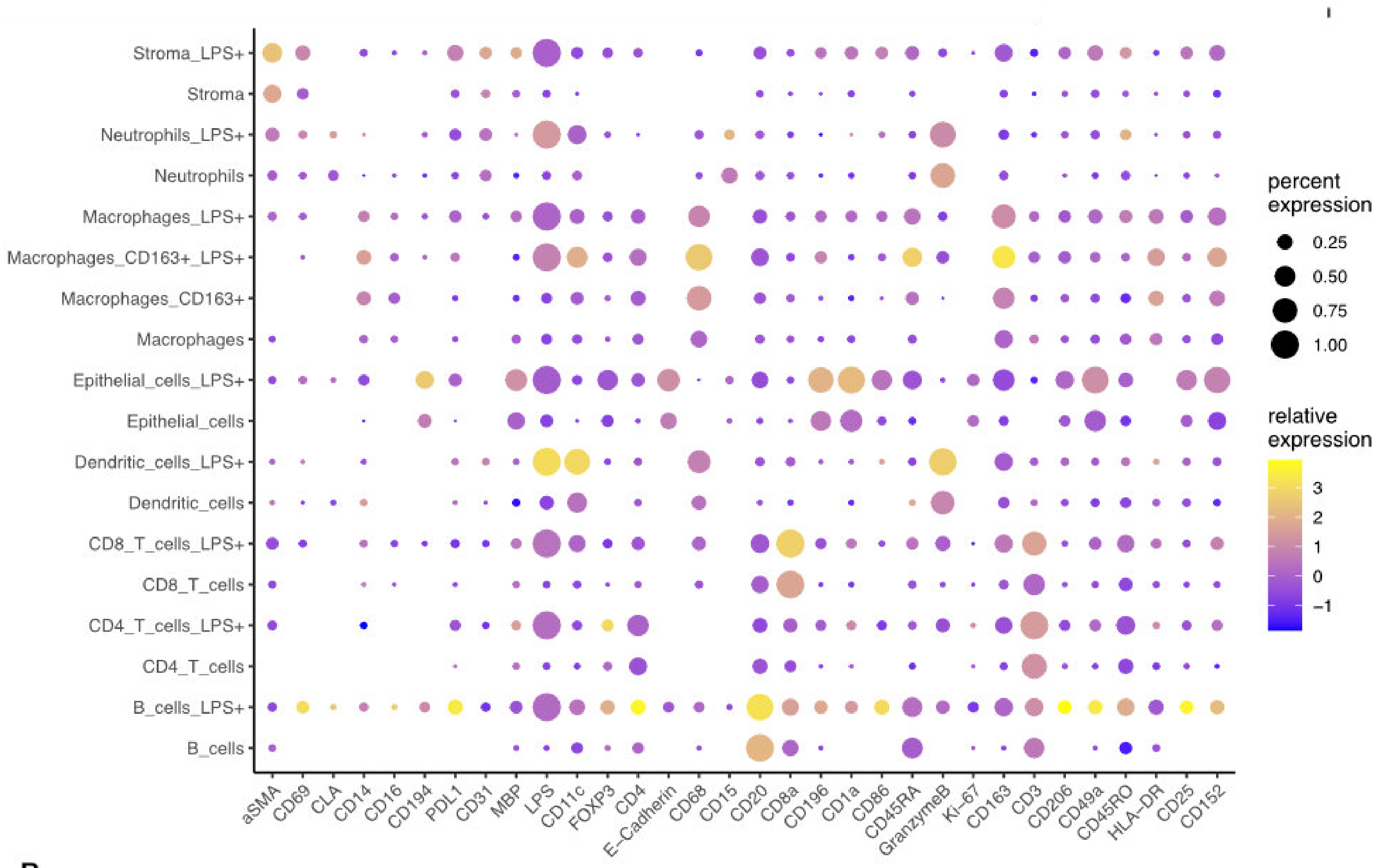

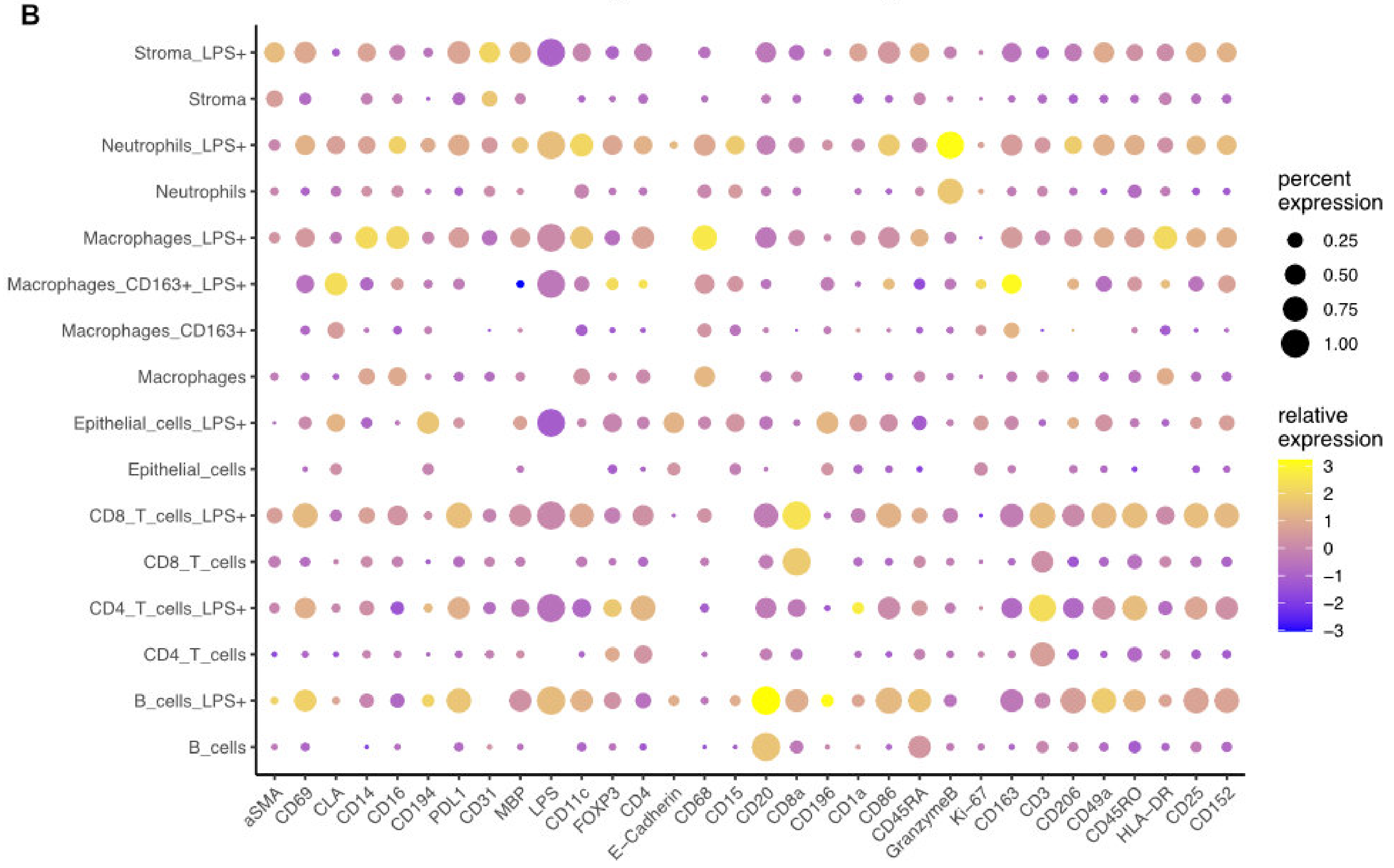

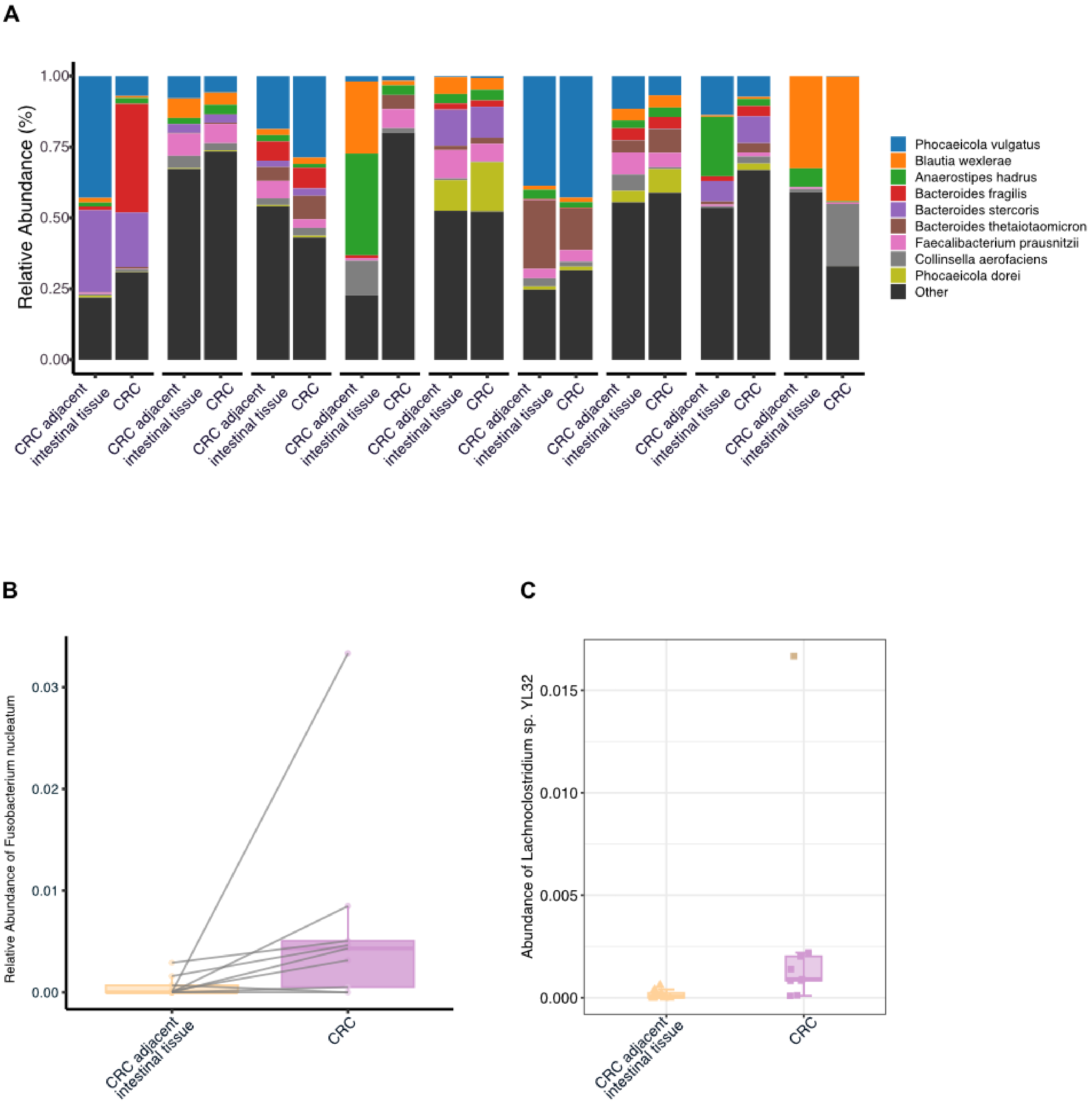
